# Response of microbial eukaryote community to the oligotrophic waters of the Gulf of Mexico: a plausible scenario for warm and stratified oceans

**DOI:** 10.1101/2023.07.26.548711

**Authors:** K. Sidón-Ceseña, M.A. Martínez-Mercado, J. Chong-Robles, Y. Ortega-Saad, V.F. Camacho-Ibar, L. Linacre, A. Lago-Lestón

**Affiliations:** Posgrado de Ciencias de la Vida, Centro de Investigación Científica y de Educación Superior de Ensenada, México; Departamento de Biotecnología Marina, Centro de Investigación Científica y de Educación Superior de Ensenada, México; Departamento de Innovación Biomédica, Centro de Investigación Científica y de Educación Superior de Ensenada, México.; Instituto de Investigaciones Oceanológicas, Universidad Autónoma de Baja California, Ensenada, México; Departamento de Oceanografía Biológica, Centro de Investigación Científica y de Educación Superior de Ensenada, México

**Keywords:** Microbial eukaryotes, 18S-rRNA metabarcoding, Gulf of Mexico, Oligotrophic ecosystem

## Abstract

In oligotrophic environments, interactions among eukaryotic microorganisms are highly complex. In the Gulf of Mexico (GoM), the Loop Current intensifies in summer and supplies the Gulf with warm and oligotrophic waters. However, mesoscale eddies within the GoM create favorable conditions for biological productivity by bringing nutrient-rich water to the subsurface layer. This study aimed to determine the structure, variability, and ecological roles of the protist in the mixed layer (ML) and deep chlorophyll maximum (DCM), representing the first V9-18S rRNA survey studying the protist community from the Southern GoM. Results revealed different assemblages between the ML and DCM. In the ML, species abundance was highly and positively correlated with temperature but negatively correlated with the nitrate concentration, whereas the opposite pattern was observed in the DCM. Alveolata represented ∼60% in both the ML and DCM, while Haptophytes and MAST dominated the ML, and Pelagophytes and Radiolarians dominated the DCM. Interestingly, *Ostreococcus* abundance increased under upwelling conditions suggesting that it may act as an indicator of the vertical nitrate flux and that picoeukaryotes respond to this instead of diatoms. Finally, our analyses revealed high levels of competition, parasitism, and predation with a high proportion of self-exclusion relationship (30%) in both depths.

## Introduction

Microbial eukaryotes (protists) show ample metabolic diversity and directly influence biogeochemical cycles in the ocean (Caron et al., 2016; Worden et al., 2015). The cell sizes of protists and their positions within food webs are directly related to their biogeochemical functions as primary producers and carbon exporters (Ward et al., 2012). Picoplankton (cell sizes of 2–3 µm) play critical roles as primary producers in oligotrophic waters (Shi et al., 2009), whereas diatoms are primarily responsible for new primary production and high rates of carbon export in upwelling regions (Bibby and Moore, 2011). Under conditions of nutrient limitation and high temperatures, the microbial loop intensifies and carbon transfer becomes less efficient due to the resulting increase in trophic complexity, with protists playing multiple roles (Azam et al., 1983) as autotrophs, heterotrophs, mixotrophs, parasites, and saprophytes and providing links to higher trophic levels (Worden et al., 2015). These adverse conditions have been exacerbated by global ocean warming, which has increased water column stratification and nutrient limitation in the mixed layer and consequently decreased primary productivity (Boyce et al., 2010; Gregg and Rousseaux, 2019).

In oceanic regions, the vertical structure of the photic zone in summer shows a strong temperature gradient that extends to the mixed layer depth and deep chlorophyll maximum (DCM). The vertical profile of the mixed layer (ML) shows almost homogeneous temperature, salinity, and density values, which are the result of turbulent, wind-induced mixing (de Boyer Montégut et al., 2004; Huang et al., 2018). Within the ML, the surface waters are warm and well-lit while nutrient availability limits primary production (Sigman et al., 2012). In the DCM, phytoplankton biomass tends to increase with the nutrient supply while light availability limits primary production (Eppley and Cullen, 1981; Sigman et al., 2012).

The Gulf of Mexico (GoM) is a semi-enclosed subtropical basin that is fed by the Loop Current (LC), which is formed by warm, saline water that enters through the Yucatan Channel and exits through the Florida Straits (Biggs, 1992; Biggs and Ressler, 2001). The oligotrophic surface water that enters the gulf via the LC does not fertilize the GoM. Instead, other conditions and mechanisms exist within the GoM that fertilize the water column and promote primary productivity. These include advective processes such as anticyclonic eddies and their edge effects, cyclonic eddies (CE), and anticyclonic-cyclonic eddy interactions (Biggs, 1992; McGillicuddy, 2016; Salmerón-García et al., 2011; Toner et al., 2003).

The three principal mesoscale structures within the GoM that are responsible for nutrient enrichment include large anticyclonic LC eddies (LCEs), CE fronts, and the semi-permanent cyclonic circulation of Campeche Bay. Large (> 200 km diameter) anticyclonic LCEs detach from the LC, travel westward toward the Tamaulipas-Veracruz shelf, and dissipate along the slope, whereas CE fronts form around the LC (Hamilton et al., 2018; Zavala-hidalgo et al., 2003). The semi-permanent cyclonic circulation of Campeche Bay operates within the southern region of the GoM (Pérez-Brunius et al., 2013). In addition, water-column nutrient enrichment can be observed throughout the year in the region of the Yucatan shelf due to upwelling, although this intensifies along with the LC in summer (Kurczyn et al., 2021). In the northern GoM, nutrient enrichment can also be observed due to fluvial discharge from the Mississippi River (Wawrik and Paul, 2004), the Grijalva-Usumacinta river system, and Veracruz-Tamaulipas platform (Martínez-López and Zavala-Hidalgo, 2009; Monreal Gomez et al., 1992).

The open ocean of the GoM is an oligotrophic ecosystem due to its high superficial temperatures in summer (28–30 °C), depleted surface nitrate concentrations, low chlorophyll concentrations (< 0.15 mg m^−3^), and temporal changes in the ML (< 40 m in summer; Damien et al., 2018; Muller-Karger et al., 2015; Pasqueron de de Fommervault et al., 2017). The principal sources of temporal chlorophyll variability in the GoM are the result of the transition from conditions of vertical water-column mixing in winter to water-column stratification in summer (Damien et al., 2018; Pasqueron de Fommervault et al., 2017; Muller-Karger et al., 2015). In addition, changes in the intensity of LC intrusion, which tend to be more intense in summer than in winter, can also affect chlorophyll concentrations (Delgado et al., 2019). Climate change predictions include a decrease in photosynthetic biomass in tropical and subtropical regions, and the greater success of smaller phytoplankton in warm, oligotrophic ocean waters (Agusti et al., 2019; Gregg and Rousseaux, 2019; Henson et al., 2021). Thus, the oligotrophic conditions of the GoM make it an excellent natural laboratory to study the possible effects of global warming.

In the oligotrophic area around Cozumel Island, Rodríguez-Gomez et al. (2022) found that vertical nutricline displacements in the subsurface layer and deep mixed layer were the principal factors that shaped the phytoplankton community, which contained a low abundance of diatoms. Studies employing flow cytometry analysis have also found that *Prochlorococcus* species dominate the ML in the oligotrophic region of the southern GoM, with the population showing the highest abundance in anticyclonic eddies in both winter and summer (Linacre et al., 2019; 2015). Furthermore, *Prochlorococcus* ecotypes that are associated with low-light conditions and may be transported by LCE dynamics have been identified near the nutricline (Linacre et al., 2019). Using fluorescent in situ hybridization coupled with tyramide signal amplification (FISH-TSA), Hernández-Becerril et al. (2012) suggested that the picoeukaryote *Micromonas pusilla* plays an important role as a primary producer and carbon exporter that is equal to those of diatoms and coccolithophorids in the Campeche region during winter.

Most of the studies performed on microbial eukaryotes in the Mexican Exclusive Economic Zone of the GoM (MEEZ) have focused on phytoplankton communities and employed microscopy analyses (Durán-Campos et al., 2017; Hernández-Becerril and Flores-Granados, 1998; Hernández-Becerril et al., 2008; Licea et al., 2016; 2004; 2011; Linacre et al., 2021; Merino-Virgilio et al., 2013; Okolodkov, 2003; Parra-Toriz et al., 2011); flow cytometry to detect *Prochlorococcus*, *Synechococcus*, heterotrophic bacteria, and picoeukaryotes (Linacre et al., 2019; 2015); and molecular analyses, such as FISH-TSA, to detect picoeukaryotes like *M. pusilla* (Hernández-Becerril et al., 2012). In contrast, metabarcoding analyses are scarce and have focused on evaluating the bacterial composition of the water column (Raggi et al., 2020) and zooplankton communities (Martinez et al., 2021). Indeed, evaluations employing metabarcoding approaches to evaluate the entire protist community in the oceanic waters of the MEEZ have not yet been conducted. Metabarcoding based on high-throughput sequencing of the V9-18S rRNA gene allows for communities to be studied with high taxonomic resolution. Through this approach, uncultivated species (Johnson and Martiny, 2015; Medlin, et al., 2011), picoplankton species that lack distinct morphological structures (Caron et al., 2012; Massana, 2011; Santi et al., 2021), and spatiotemporal differences (Santoferrara et al., 2020) may be identified, resulting in a more accurate picture of microbial communities.

In this study, we used a metabarcoding approach to evaluate potential differences in the protist community between the ML and DCM of the GoM during warm and intensely stratified summer conditions. To this end, we characterized the protist assemblages of the ML and DCM, evaluated which environmental factors shape protist community structure in the ML and DCM, and assessed how protist-protist relationships could shape the community. We believe that the results of this study will contribute to a better understanding of how protist communities may respond to changes in highly stratified environments, thus establishing a baseline for future studies of important oligotrophic ecosystems within the context of global climate change.

## Experimental procedures

### Sample collection

Ninety-two samples were taken from the euphotic zone of the deep oceanic region of the southern GoM during four oceanographic cruises: XIXIMI-4 (August 27 to September 16, 2015), XIXIMI-5 (June 10–25, 2016), XIXIMI-6 (August 15 to September 8, 2017), and XIXIMI-7 (May 9 to June 2, 2019) onboard the R/V Justo Sierra (UNAM). Seawater was sampled in 44 oceanic stations at two different depths: the ML (5–10 m) and DCM (85 m). Some stations were included in multiple cruises (Fig. 1). Seawater was collected using 20-L Niskin bottles mounted to a 12-bottle SBE32 carousel (Sea-Bird Scientific, Bellevue, USA), and the hydrographic data were collected using a SBE 911plus CTD (Sea-Bird Scientific) equipped with a Seapoint fluorometer (Seapoint Sensors Inc., Exeter, USA) and an SBE43 dissolved oxygen sensor (Sea-Bird Scientific). For each sample, 6 L of seawater was prefiltered through 200-µm Nitex mesh to limit the presence of multicellular eukaryotes, and then split into two 3-L portions for vacuum filtration using Whatman® Nuclepore™ track-etched membranes (47-mm diameter, 0.8-µm pore size; Millipore, Burlington, USA) to collect the 0.8–200 µm fraction. After filtration, all samples were immediately frozen in liquid nitrogen and stored at −80 ᵒC until further processing.

**Figure 1.**
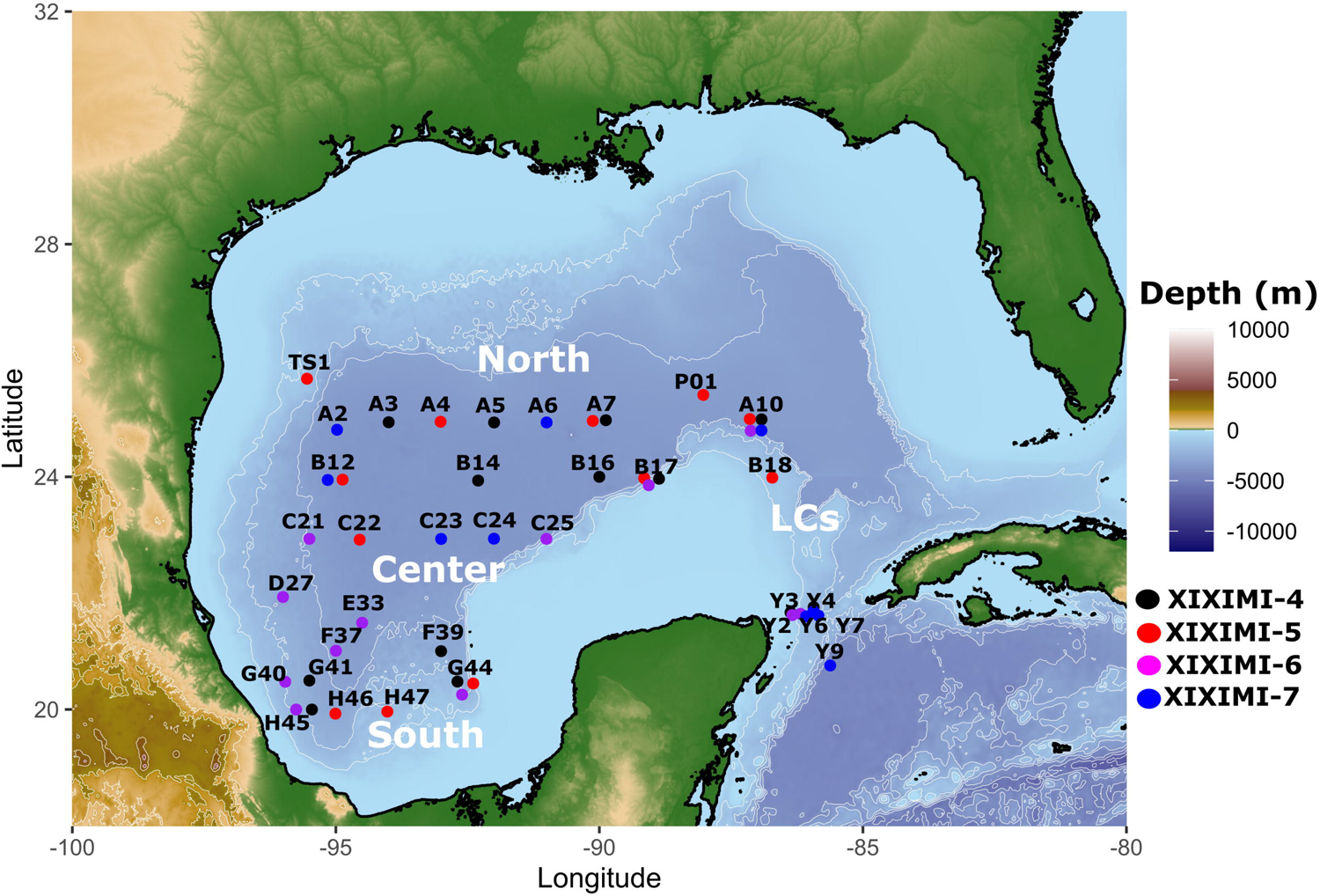
Geographic locations of the stations in which microeukaryotes were sampled during the XIXIMI-4 (2015), XIXIMI-5 (2016), XIXIMI-6 (2017), and XIXIMI-7 (2019) oceanographic cruises, in the exclusive economic zone of Mexico (EEZ) in the deep-water region (bottom depth > 1000 m) of the Gulf of Mexico (GoM). The sampling area was divided into the Loop Current (LC; stations within the LC and Yucatan Channel), northern (north of the MEEZ; 25–23° N), central (center of the GoM; 22–21° N), and southern (Bay of Campeche; 20–19 ° N) regions.

### Environmental conditions

The ML was estimated using the individual density profile method of Huang et al. (2018), which employs a quality index (QI) adapted from Lorbacher et al. (2006). Additionally, we determined the Brunt-väisäla frequency (N^2^) as a proxy of water column stability (Millard et al., 1990).

In order to organize the stations according to mesoscale influence, the vertical displacement of the subsurface 25.5 kg m^−3^ isopycnal was used as a proxy of the nutricline (Pasqueron de Fommervault et al., 2017). For this, the average depth of the 25.5 kg m^−3^ isopycnal for all CTD casts was determined by cruise. Stations were classified as being under anticyclonic eddy influence (downwelling conditions) when the depth of the 25.5 kg m^−3^ isopycnal was at least one standard deviation above the overall average. Similarly, stations under the influence of CE (upwelling conditions) were identified when the depth of the 25.5 kg m^−3^ isopycnal was at least one standard deviation below the overall average.

An analysis of the nutrients present at the ML and DCM depths was conducted with an automated AA3-HR nutrient analyzer (Seal Analytical, Mequon, USA) following the guidelines described in the GO-SHIP Repeat Hydrography Manual (Hydes et al., 2010). Nitrate+nitrite (hereinafter referred to as nitrate) and silicic acid concentrations were determined according to the method of Armstrong et al. (1967) with modifications. Reference materials for nutrients in seawater [lot CC and lot CD developed by The General Environmental Technos Co., Ltd. (Kanso Technos, Osaka, Japan; Aoyama & Hydes, 2010) were repeatedly analyzed during runs to evaluate accuracy and precision. The limits of detection for the nitrate and silicic acid concentrations were 0.02 and 0.04 µmol kg^−1^, respectively.

### DNA extraction, PCR amplification, and library preparation

Total DNA was extracted from the membranes using the DNeasy Power Water Kit (Qiagen, Hilden, Germany) with the addition of 200 µL of phenol:chloroform:isoamyl alcohol (25:24:1; Sigma-Aldrich, St. Louis, USA) during the cell lysis process to increase DNA yield. Libraries were generated via one-step PCR using primers containing the specific oligos 1389F (5’-GCCTCCCTCGCGCCATCAG −3’) and 1510R (5’-GCCTTGCCAGCCCGCTCAG −3’; Amaral-Zettler et al., 2009) to amplify the V9 region of the 18S rRNA gene. The Illumina adapters and dual-index barcodes were constructed according to the method of Kozich et al. (2013). The PCR reactions were performed in duplicate for each sample using 10 ng template DNA, 1X Mix MyTaq™ (Bioline, London, UK), 0.4 uM of each of the forward and reverse primers, and 0.6 uM of MyTaq^TM^ DNA polymerase (Bioline). The reaction was carried out in a total volume of 20 µl with the following thermal conditions: an initial denaturation of 95 °C for 5 min, followed by 32 cycles of 95 °C for 20 s, 55 °C for 15 s, 72 °C for 5 min, and 72 °C for 7 min. We included a negative PCR control, which did not amplify. The PCR products were cleaned up and normalized, and fragment size selection was conducted in SequalPrep^TM^ (Thermo Fisher Scientific, Waltham, USA). The normalized PCR products were quantified using a Qubit 3.0 Fluorometer and the dsDNA HS (high sensitivity) Assay Kit (Thermo Fisher Scientific). Finally, the libraries were pooled to equimolar concentrations and paired-end sequenced (2 x 151) using the MiSeq Reagent kit V2® on an Illumina MiSeq (San Diego, USA) platform at the Centro de Investigación Científica y de Educación Superior de Ensenada (CICESE).

### Sequence data processing and taxonomic assignment

Amplicon sequencing was conducted in two sequencing runs, and the data from each run were processed independently. Primers and adapters were trimmed, and reads with homopolymers containing more than 10 bases and a minimum length of 70 bp were discarded using Cutadapt v. 2.5 (Martin, 2011). The trimmed sequences were processed with the DADA2 v. 1.8 pipeline (Callahan et al., 2016) and quality filtered and denoised with the parameters *minLen* 80, *maxEE* 2, and *truncQ* 10 and the merging parameters of *minOver* 10 and *maxMM* 0 in R v. 3.5.0 (R Core Team, 2017). The amplicon sequence variants (ASVs) obtained from both datasets were merged into a single sequence table, and the chimeras were removed using the consensus method. Taxonomic assignments were performed against the Protist Ribosomal Reference database (PR^2^) v. 4.12 (Guillou et al., 2012) with the naive Bayesian classifier method using the function *assignTaxonomy* (bootstrap > 80) of DADA2. The ASVs affiliated to the Opisthokonta Supergroup and the ASVs not assigned (NA) at any division were removed from further analysis. The results were formatted as *phyloseq objects* (McMurdie and Holmes, 2013) for ecological analyses. Alpha diversity indices (Shannon and Richness) were calculated via the R package ‘*microbiome*’ (Lahti L, et al. 2019) using the *global* function.

### Ecological and Statistical Analyses

For the beta diversity analysis, the read counts were normalized with a centered log ratio (clr) transformation to account for the different number of sequences due to the compositional nature of the data (Gloor et al., 2017). The environmental parameters were normalized by Z-score for multivariate analyses. A non-metric multidimensional scaling (NMDS) analysis was performed based on Aitchinson distances calculated from Euclidean distances with the ‘*vegan*’ package v. 2.5 (Oksanen et al., 2020). An analysis of similarity (ANOSIM; 999 permutations and Aitchinson distance matrix) was conducted with the ‘*vegan*’ package to detect significant differences at the community level between groups of samples. A permutational analysis of variance (PERMANOVA; 999 permutations and Aitchinson distance matrix) was performed using the *adonis* function to detect significant differences between regions, cruises, and nutricline conditions (upwelling, downwelling, or neutral). We used the distance-based redundancy analysis (db-RDA) to determine which environmental factors were the most important in explaining the variation in microeukaryote community composition (Legendre and Anderson, 1999). Environmental variables were selected when the variance inflation factor (VIF) was less than four. This parameter resulted in the selection of temperature (°C), nitrate (µmol/kg), oxygen (µmol/kg), latitude, absolute salinity, silicic acid (Si), and dissolved organic carbon (DOC; µmol/kg). We evaluated the statistical significance of the db-RDA model and analyzed each environmental variable that significantly contributed to the variation in community composition with an ANOVA-like permutation test (Legendre et al., 2011).

The correlations among the abundances of the 35 most abundant taxa and environmental variables were evaluated with an sPLS (sparse Partial Least Squares) regression with the *spls* () function of the ‘*Mixomics*’ package in R (Rohart et al., 2017). This technique is not limited to uncorrelated variables and can use noisy, collinear (correlated), or missing variables and can predict one group of variables from another group of variables (Lê Cao et al., 2008). In this study, the analysis predicted the abundance of the principal microeukaryote taxa from the environmental variables.

Wilcoxon signed-rank tests were performed to evaluate the differences in abundance among the 35 principal microeukaryotes (excluding Alveolata) and the vertical distribution of the subsurface 25.5 kg m^−3^ isopycnal and to evaluate changes in mean abundance due to upwelling, downwelling, and neutral conditions. Furthermore, we used Deseq2 (differential abundance analysis) based on a negative binomial distribution (Love et al., 2014) to evaluate if the abundance of ASVs differed between the ML and DCM.

The covariance network was analyzed using SPIEC-EASI (sparse inverse covariance estimation for ecological association inference) v. 1.1.0 (Kurtz et al., 2015) with the “mb (Meinshausen-Buhlmann) neighborhood method with 50 repetitions and ASVs (taxonomy grouped to the lowest taxonomic rank). The keystone species were detected with a degree value > 10 (i.e., the number of associations/edges connected per node/ASV) and betweenness > 1000 (the total number of shortest paths from all nodes to all other nodes passing through that node). Finally, positive and negative associations/edges (> 0.02) were filtered from the network with the top 80 ASVs most prevalent at each depth. The network statistic was conducted with the R package ‘*igraph*’ (Csárdi and Nepusz, 2006), and the network associations were visualized with chord diagrams using the R package ‘*Circlize*’ (Gu et al., 2014).

## Results and Discussion

We conducted a V9-18S rDNA-based metabarcoding survey to evaluate the diversity of the protist community of the oligotrophic region of the GoM during the warm season to determine how environmental and biological factors shape this community. We found notable differences between the two water-column layers evaluated in this study. The ML was dominated by mixotrophic and heterotrophic picoeukaryotes due to high vertical thermal stratification, whereas mixotrophic and photosynthetic picoeukaryotes were dominant in the DCM given the broad variation in nutrient concentrations and low light conditions of this depth. Furthermore, a high degree of negative associations of parasitism and predation may influence the protist assemblages in both the ML and DCM.

### Environmental conditions of the photic zone

At the time of sampling, the water column exhibited high stratification, which was evident by the shallow values of the average ML (XIXIMI-4 = 31 m, XIXIMI-5 = 22 m, XIXIMI-6 = 28 m, and XIXIMI-7 = 40 m). In summer, ML values lower than 40 m (Fig. 2A) have been found in the gulf with long-trend analyses (Muller-Karger et al., 2015; Portela et al., 2018). The mixed layer was characterized by high temperatures (27.2 to 30.6 °C; Fig. 2B), a wide range of salinity values (34.4 to 36.6; Fig. 2C), and low nitrate concentrations (0.54 µmol kg^−1^ to the detection limit of 0.02 µmol kg^−1^; Fig. 2d). In contrast, silicic acid (silicate) was only high (> 2 µM) in the southern region (Fig. 2E). Previous studies have reported low surface chlorophyll (Chla) concentrations during oligotrophic conditions in the ML, particularly during summer (Damien et al., 2018; Pasqueron de Fommervault et al., 2017). In this study, the DCM was characterized by a low average fluorescence value when it was deeper (0.42 RFU; relative fluorescence units) and a high average fluorescence value when it was shallower (1.34 RFU; Fig. S1). The temperature of the DCM ranged from 20.5 to 27.7 °C, while salinity ranged from 36 to 36.6, and the maximum nitrate concentration ranged from 5.2 µmol kg^−1^ to below the detection limit (Fig. 2B-E). The nutricline (Fig. 2F, Fig. S2), was shallower during XIXIMI-4 (87 m) than during XIXIMI-5 (88.7 m), XIXIMI-6 (120 m), or XIXIMI-7 (112 m). Notably, station PO1 was sampled during XIXIMI-5. This station was located at the core of the recently detached LCE Poseidon (Horizon Marine group, www.horizonmarine.com/loop-current-eddies) and exhibited the deepest nutricline depth (255 m) of the data set. Other stations with deep nutricline values included A10 (XIXIMI-5 = 167 m; XIXIMI-6 = 217 m; and XIXIMI-7 = 238 m) located in the LC and Y7 (XIXIMI-7 = 157 m) and Y9 (XIXIMI-7 = 244 m) located in Caribbean waters (Fig. 1).

**Figure 2.**
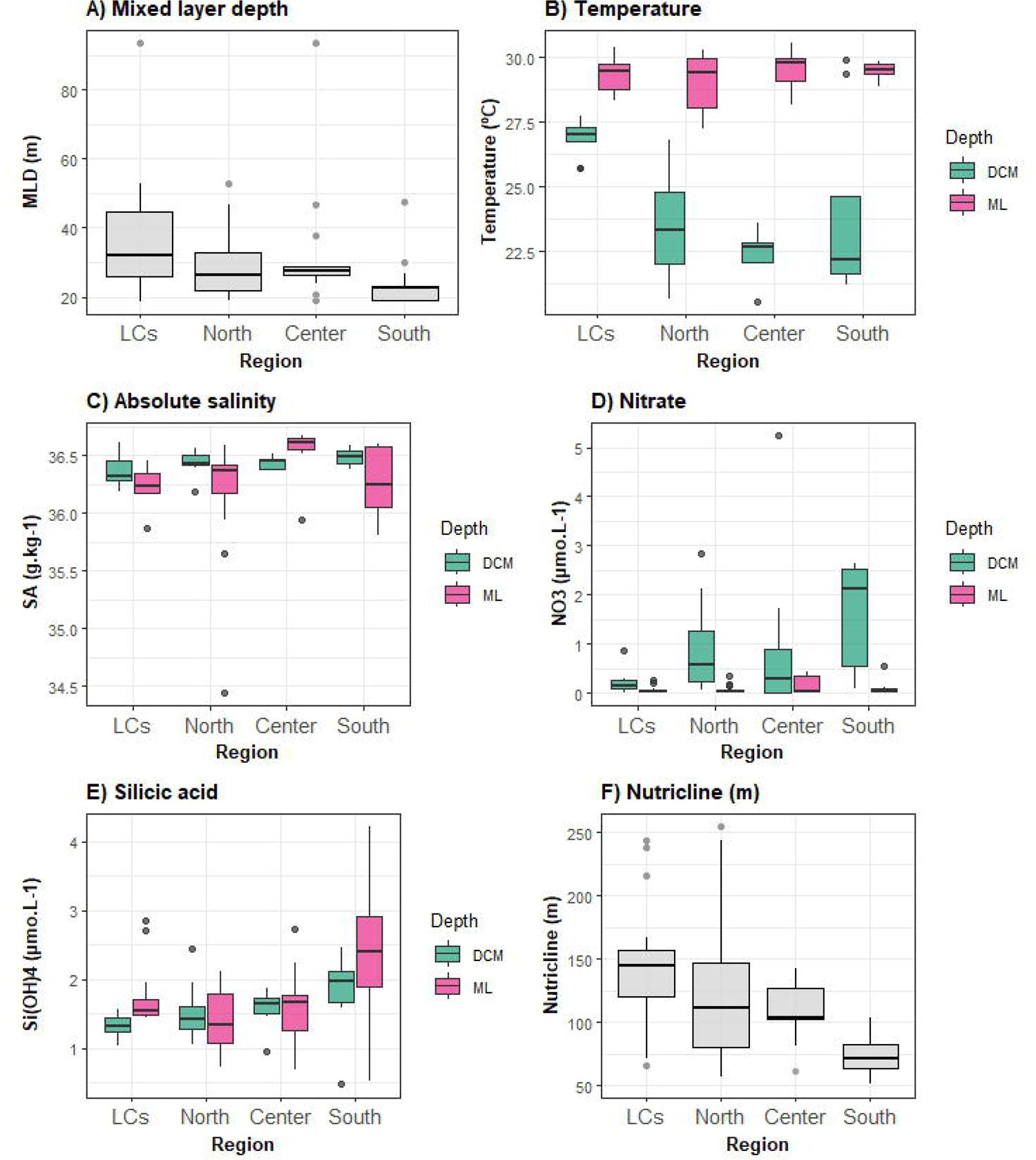
Environmental conditions of the surface water at the two sample depths: the mixed layer (ML) and deep chlorophyll maximum (DCM). Samples were grouped as Loop Current (LC), northern, central, and southern stations and mean values were calculated by group for each depth. (A) MLD (m), (B) temperature (°C), (C) absolute salinity (g kg^−1^), (D) nitrate (µmol kg^−1^), (E) silicic acid (µmol kg^−1^), and (F) nutricline (m, distribution depth of the isopycnal 25.5). The green and pink colors represent the DCM and ML, respectively (Figures B, C, D, E, F).

From a regional point of view, the area associated with the CE of the Bay of Campeche was the most stratified (ML = 21.4 m, nutricline = 73 m; Fig. 2A, Fig. 2F), whereas the stations inside the LC and the northern and central regions of the gulf were the least stratified (average ML of 40 m, average nutricline depth of 167 m, Fig. 2A, Fig. 2F). Moreover, it has been reported that the effects of the CE in the Bay of Campeche were not observed in the ML during XIXIMI-5 due to high temperatures in the ML despite shallow isopycnals (25.5 isopycnal; Lee-Sánchez et al., 2022). The average nitrate concentration in the southern gulf only exhibited low nitrate values in the ML (0.1 µmol kg^−1^) with higher values observed in the DCM (1.2 µmol kg^−1^; Fig. 2D). In contrast, the average nitrate concentration in LC stations was low (0.1 µmol kg^−1^) at both depths (ML and DCM; Fig. 2D). Overall, the strong stratification of oceanic waters, which is characteristic of summer conditions, resulted in a clear difference between the conditions of the ML and DCM.

### Protist community composition and structure in the ML and DCM

A total of 4,524,956 reads were generated from the 92 samples, with an average of 49,184 reads per sample. After sequence processing, 4,889 ASVs were obtained with an average of 453 ASVs per sample (max 871, min 81). In both depths, Alveolata was the most abundant taxa, with a total of 1,593,897 reads (69.75%) in the ML and 1,212,367 reads (54.13%) in the DCM (Fig. 3A-B). In the ML, Stramenopiles represented 14.49% of all reads, followed by Hacrobia (10.2%) and Rhizaria (3.86%). In the DCM, Alveolata was followed by Rhizaria (21.59%) and Stramenopiles (13.49%) as the most abundant taxa while the relative abundance of Hacrobia decreased to 6.88% (Fig. 3A-B). Although Alveolata dominated the communities at both depths, the presence of Rhizaria was also notable, as its abundance transitioned from low in the ML to the second most-abundant taxa in the DCM.

**Figure 3.**
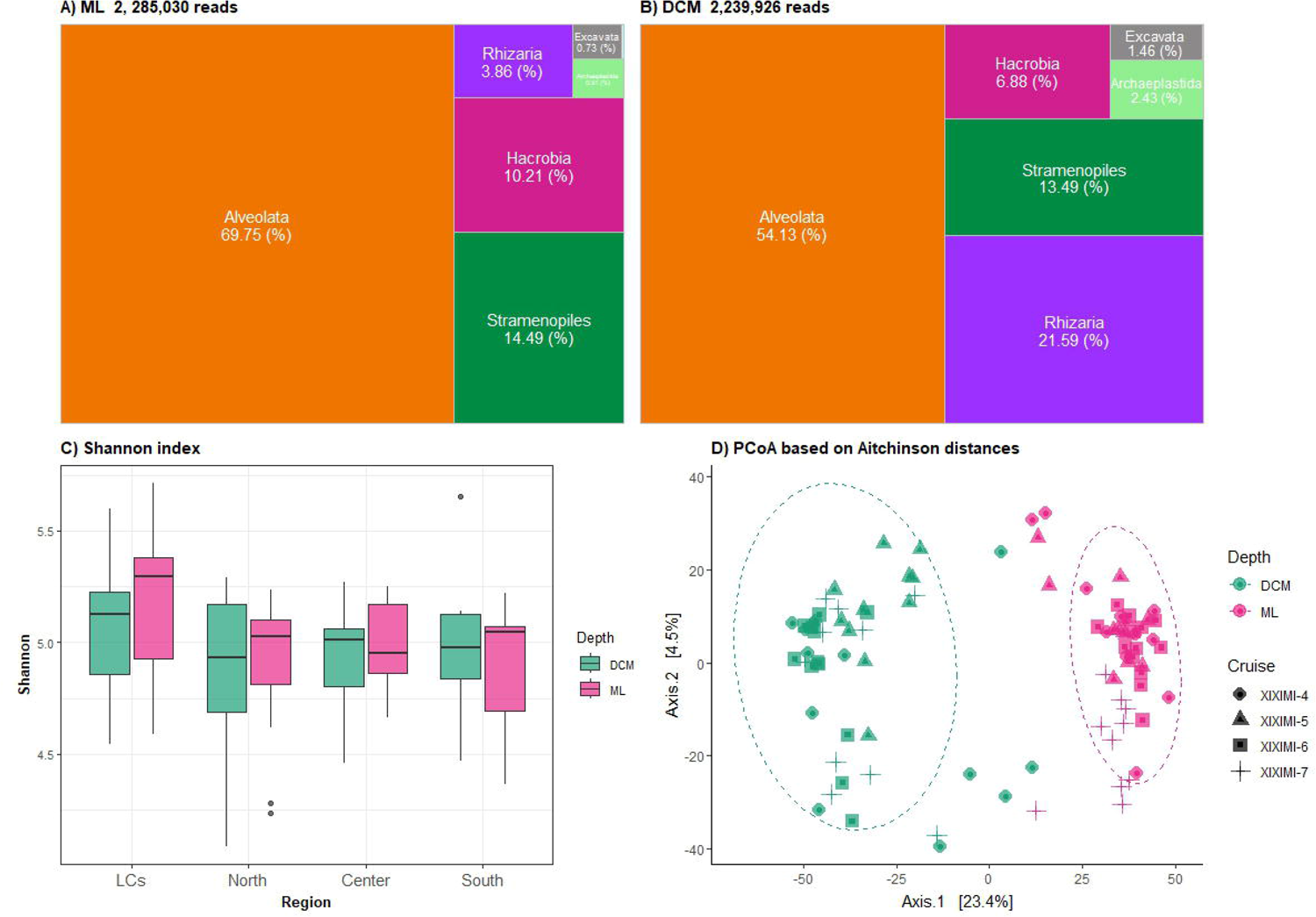
Alpha and beta diversity patterns. The treemaps show the distribution of the reads from the taxonomic level of the supergroup: (A) Alveolata, Stramenopiles, Hacrobia, Rhizaria, Archaeplastida, and Excavata in the mixed layer depth (ML) and (B) deep chlorophyll maximum (DCM). (C) Shannon index by region [Loop Current (LC), northern, central, and southern]. (D) Non-metric multidimensional scaling (NMDS) based on Aitchinson distances (stress = 0.12) from the amplicon sequence variants (ASVs) of the protist community from the ML (pink) and DCM (green). The shape indicates the cruise.

The Shannon diversity index values ranged from 4.09 to 5.7 for all samples (Fig. 3C). The value of this index was significantly different by cruise (p = 0.000). In contrast, no significant differences were found in Shannon diversity when comparing values between the ML and DCM (p = 0.80) or between nutricline depths (p = 0.36). Of note, the samples from the LC were the most diverse (Fig. 3C), with significant differences between LC stations and those in the northern (p = 0.006) and southern (p = 0.043) regions of the gulf. These differences may be associated with the high temperature and low nitrate values observed in the ML and DCM in the LC (Fig. 2). Temperature is the principal driver behind the notable increase in protist community diversity from the poles to subtropical regions (Ibarbalz et al., 2019), although the oceanographic conditions in the surface layers and very low primary productivity also influence diversity (Raes et al., 2018). This agrees with the diversity patterns that have recently been reported for the Sargasso Sea, with high diversity being observed during warm, oligotrophic conditions (Blanco-Bercial et al., 2022) and in the water column of the Gulf stream (Countway et al., 2006).

The NMDS analysis indicated that the protist community is clustered by depth (Fig. 3D), and the samples from the ML show greater similarity to each other than those of the DCM, which showed more variability among stations. The composition of the protist community was significantly different and dissimilar by depth (ANOSIM p = 0.001, R = 0.93; PERMANOVA R^2^ = 0.38, p = 0.001). These results agree with those of other studies conducted in different oceanic regions of the Atlantic and Pacific oceans (Countway et al., 2007; Brisbin et al., 2020; Ollison et al., 2021) and those of global scale analyses (de Vargas et al., 2015; Ibarbalz et al., 2019), which shows that different environmental conditions based on depth lead to differences in community structure. The opposing light and nutrient patterns between the ML and DCM led to dissimilarities between these two depths. However, when community structure was constrained to environmental conditions, the RDA model explained a low percentage of the variation in the ML and DCM (Fig. 4). In the ML, the RDA model (dbRDA, p = 0.001) indicated that silicic acid, temperature, oxygen, latitude, and salinity significantly influenced the variability in community structure (Table 1), although these only explained 15.8% of the variability (Fig. 4A). Compared to what was observed in the DCM, the RDA model was significant (dbRDA, p = 0.001); however, only nitrate and temperature significantly influenced the variability in community structure (p = 0.001, Table 1) and only explained 10.7% of the variability (Fig. 4B).

**Figure 4.**
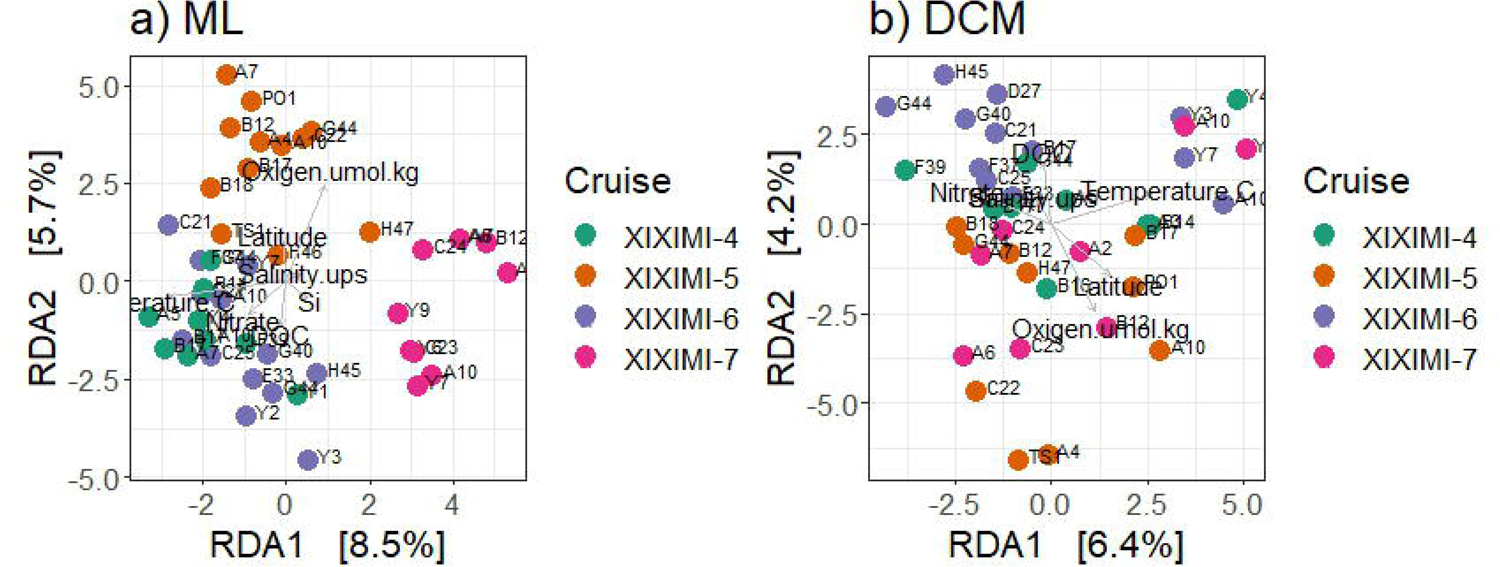
Redundancy analysis (db RDA) of the community composition of protists from the (A) mixed layer depth (ML) and (B) deep chlorophyll maximum (DCM) based on Aitchinson distances. The dot color indicates the cruise. Vectors represent the selected environmental variables. The lengths and directions of the vectors represent the correlations of the environmental variables of temperature, nitrate, oxigen, salinity, disolved organic carbon (DOC), silicic acid, and latitude.

**Table 1.**
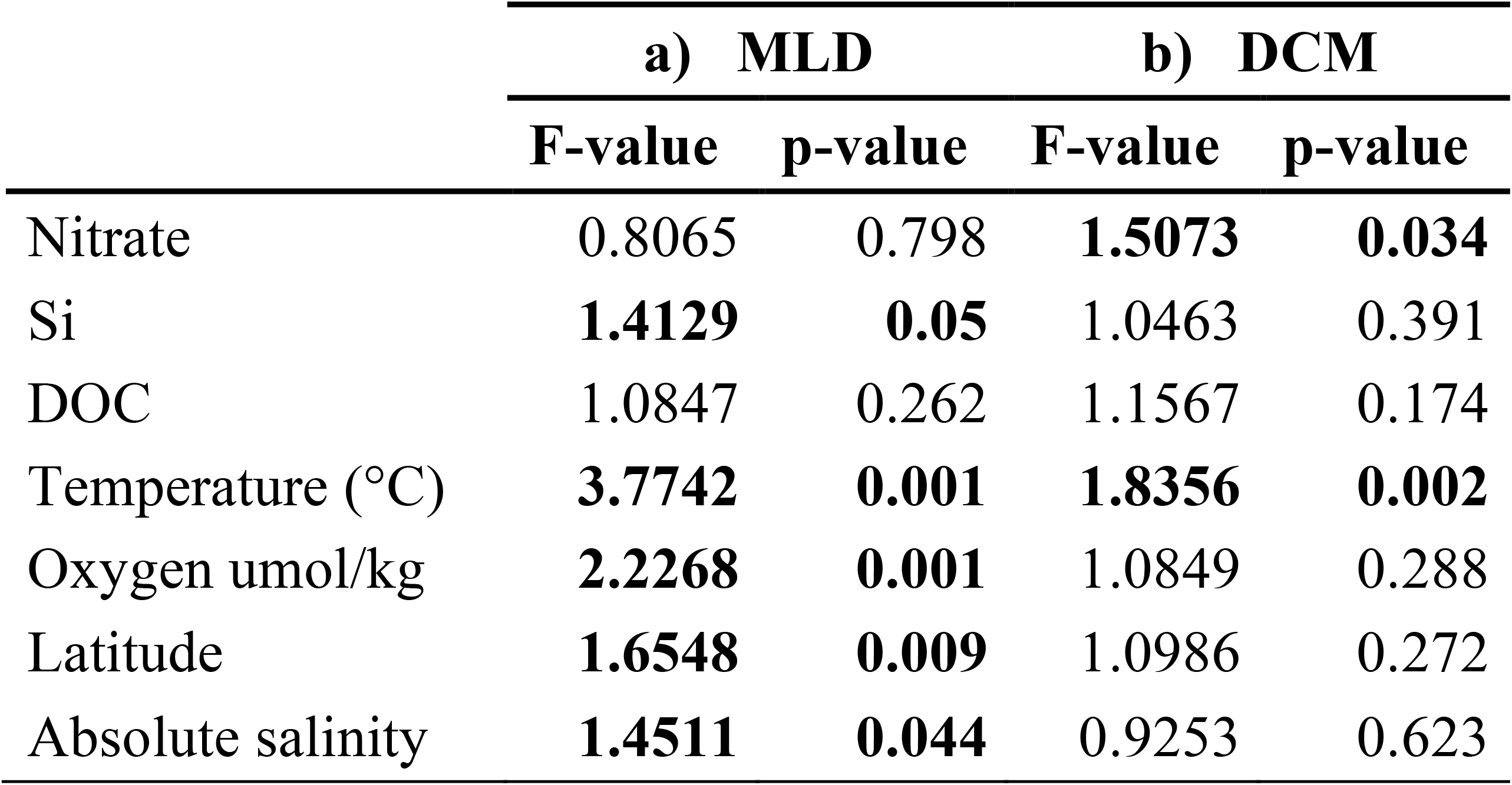
ANOVA-like permutation test for RDA by term with 999 permutations for (A) Mixed layer depth, and (B) Deep Chlorophyll maximum.

|  | a) MLD |  | b) DCM |  |
| --- | --- | --- | --- | --- |
|  | F-value | p-value | F-value | p-value |
| Nitrate | 0.8065 | 0.798 | **1.5073** | **0.034** |
| Si | **1.4129** | **0.05** | 1.0463 | 0.391 |
| DOC | 1.0847 | 0.262 | 1.1567 | 0.174 |
| Temperature (°C) | **3.7742** | **0.001** | **1.8356** | **0.002** |
| Oxygen umol/kg | **2.2268** | **0.001** | 1.0849 | 0.288 |
| Latitude | **1.6548** | **0.009** | 1.0986 | 0.272 |
| Absolute salinity | **1.4511** | **0.044** | 0.9253 | 0.623 |

The protist community was significantly different between LC stations (Fig. 1) when compared to those within the gulf (southern, central, and northern GoM, Fig. 1) within the ML (PERMANOVA, R^2^ = 18%, p = 0.001) but not in the DCM (Betadisper, p = 0.016). The pairwise comparison also revealed significant differences in LC stations among regions within the gulf, although only in the ML (Table S1–S2). These results indicate that the protist community of the ML is different compared to that in the gulf.

The categorical variable of cruise was significant only for the ML (PERMANOVA, R^2^ = 18.3%, p = 0.001) but not for the DCM (Betadisper, p = 0.035). The PERMANOVA pairwise comparison among cruises only showed a significant difference with regard to XIXIMI-5 (Table S1). These results may indicate that some seasonality was present, although we only sampled during late spring and summer. The shift from vertical mixing to stratified conditions between winter and summer (Damien et al., 2018) and the annual variability of LC intrusion (Delgado et al., 2019) are factors that promote temporal variability, which have been shown to affect the high seasonality of the protist community in the northern GoM (Brannock et al., 2016) and Sargasso Sea (Blanco-Bercial et al., 2022) and are known to be associated with environmental changes at the surface. Moreover, station PO1 should be considered to reflect extreme conditions, which was evident by this station showing the highest variability of XIXIMI-5. Notably, the deepest nutricline was present at station PO1, which was located within the recently detached LCE.

Finally, the nutricline was significantly different in the DCM (PERMANOVA R^2^ = 11%, p = 0.001) but not in the ML (PERMANOVA, R^2^ = 6%, p = 0.542). The pairwise comparison between downwelling and upwelling was significant (p = 0.003) in the DCM (Table S2). This indicates that the vertical movement of the nutricline influences the protist community only in the DCM but not in the ML. However, the low percentage of explained variation in the RDA analysis in the ML and DCM (Fig. 4) suggests that other factors, such as biotic relationships, also shape the structure of the protist community during the warm season in the GoM.

### Alveolata dominates the ML and DCM

Alveolata was the most abundant group in both layers. In particular, the Dinophyceae class was the most abundant (45–51%) and diverse (1,313 ASVs) taxa at both depths and was mainly represented by *Gymnodinium* spp. (Fig. 5A, Fig. S3), as has been observed for the subtropical and oceanic region of the GoM based on V9-18S rDNA data (Brannock, et al., 2016; Le Bescot et al., 2016). This abundance is often overestimated due to the high copy number of the rRNA gene; however, previous studies have reported that the sequence abundances of Alveolata correlate with its biovolume and biomass (Gong et al., 2013; Liu et al., 2021; Zhu et al., 2005). Accordingly, measurements based on microscopy in the oceanic region of the GoM have shown that *Gymnodinium* spp. with cell sizes < 20 µm comprise up to 55% of the biomass in summer (Linacre et al., 2021). This heterotrophic genus is associated with the mortality of phytoplankton grazers in the northern GoM (Strom and Strom, 1996) and in other oligotrophic regions such as the subtropical Atlantic (Quevedo and Anadón, 2001).

**Figure 5.**
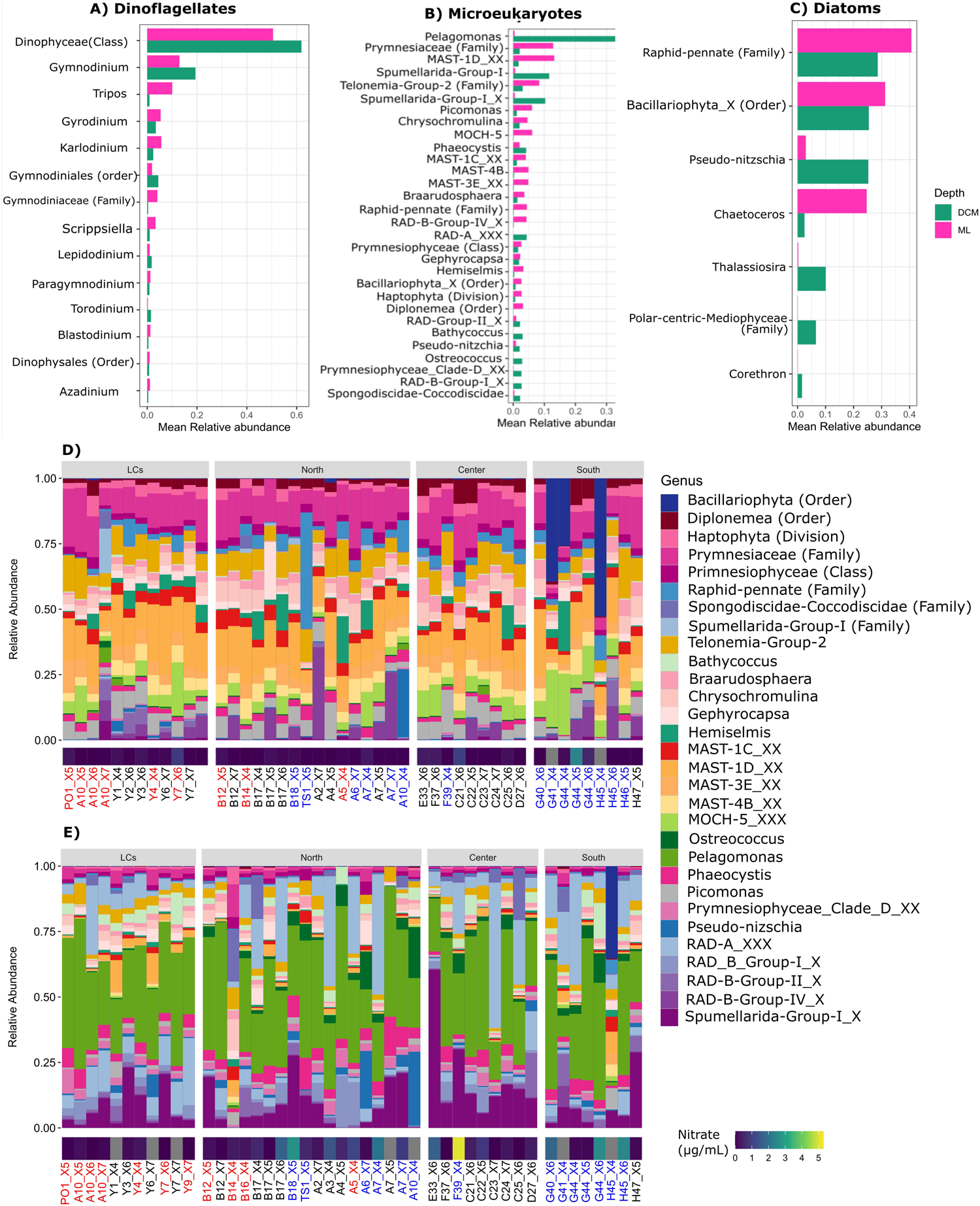
The panels of the top represent the mean abundance of (A) Dinoflagellates, (B) Microeukaryotes, and (C) Diatoms. Relative abundance of the principal protists excluding Alveolata from (D) the mixed layer (ML) and (E) the deep chlorophyll maximum (DCM). The sampling stations were ordered by region: LCs (Loop Current stations within the LC and the Yucatan Channel), northern [region to the north of the Exclusive Economic Zone of Mexico (MEEZ)], central [center of the Gulf of Mexico (GoM)], and southern (Bay of Campeche). The heatmap at the bottom of each panel represents nitrate concentration, and the gray color indicates an absence of data. The stations labeled in blue and red indicate that upwelling and downwelling conditions, respectively, were present at the time of sampling based on the vertical displacement of 25.5 kg m^−3^ isopycnal (See Fig. S2).

Syndiniales, Ciliophora, and Apicomplexa showed low abundances compared to those of dinoflagellates (Fig. S3). Syndiniales, which form part of the picoplankton fraction (2–3 µm), are parasites that infect copepods and other protists (Anderson and Harvey, 2020; Guillou et al., 2008; Zamora-Terol et al., 2021), and their abundance usually increases according to the depth of the active community below the euphotic zone (rRNA; Giner et al., 2019; Ollison et al., 2022) in oceanic regions. The co-occurrence analysis revealed a keystone (Table 2) species from this order, ASV_01050: Dino-Group-II-Clade-7_X, which has been associated with the photic zone (Guillou et al., 2008). In the northern GoM, the abundance of Syndiniales is positively correlated with copepod abundance, which are known hosts for this zooplankton group (Brannock et al., 2016). In addition, the Calanidae family was highly abundant in the water column during the oceanographic campaign XIXIMI-5 (Martinez et al., 2021). Despite their low abundance (Fig. S3), Syndiniales play a central role in the ecosystem. This highly diverse and prevalent group (270 ASV) exerts a strong influence over biogeochemical cycles by incorporating dissolved organic matter (DOM) and particulate organic matter (POM) into the microbial loop through host lysis (Anderson and Harvey, 2020; Salomon et al., 2009). Our data suggest that Syndiniales, in addition to Dinoflagellates, could shape the structure of the protist community in both the ML and DCM, exerting top-down control by parasitizing and grazing heterotrophic and mixotrophic dinoflagellates.

**Table 2.**
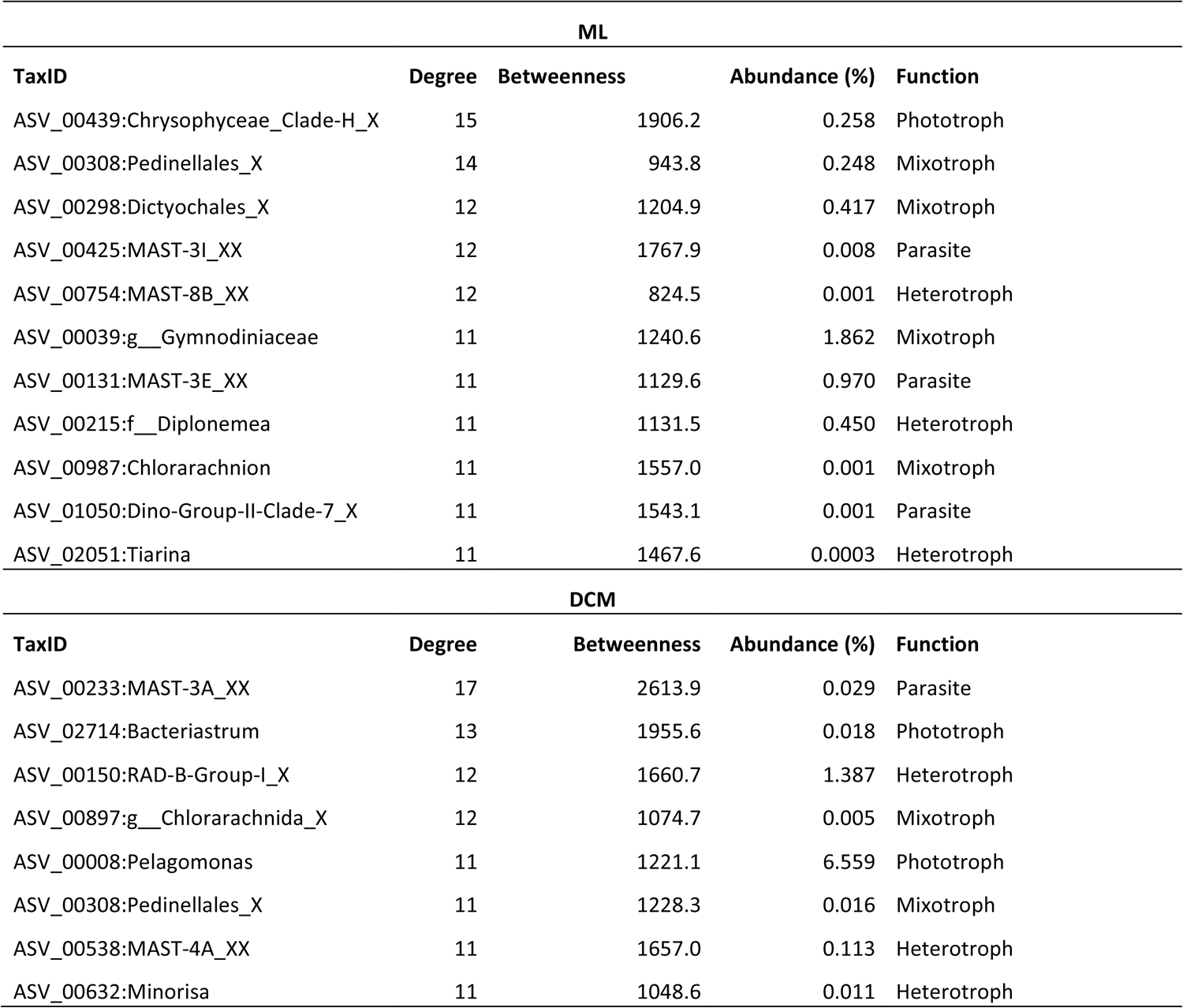
Hubspecies. List of species with high degree and high betweenness, abundance and functional annotation according to Ramond et al. (2019).

| ML |  |  |  |  |
| --- | --- | --- | --- | --- |
| TaxID | Degree | Betweenness | Abundance (%) | Function |
| ASV_00439:Chrysophyceae_Clade-H_X | 15 | 1906.2 | 0.258 | Phototroph |
| ASV_00308:Pedinellales_X | 14 | 943.8 | 0.248 | Mixotroph |
| ASV_00298:Dictyochales_X | 12 | 1204.9 | 0.417 | Mixotroph |
| ASV_00425:MAST-3I_XX | 12 | 1767.9 | 0.008 | Parasite |
| ASV_00754:MAST-8B_XX | 12 | 824.5 | 0.001 | Heterotroph |
| ASV_00039:g__Gymnodiniaceae | 11 | 1240.6 | 1.862 | Mixotroph |
| ASV_00131:MAST-3E_XX | 11 | 1129.6 | 0.970 | Parasite |
| ASV_00215:f__Diplonemea | 11 | 1131.5 | 0.450 | Heterotroph |
| ASV_00987:Chlorarachnion | 11 | 1557.0 | 0.001 | Mixotroph |
| ASV_01050:Dino-Group-II-Clade-7_X | 11 | 1543.1 | 0.001 | Parasite |
| ASV_02051:Tiarina | 11 | 1467.6 | 0.0003 | Heterotroph |
| DCM |  |  |  |  |
| TaxID | Degree | Betweenness | Abundance (%) | Function |
| ASV_00233:MAST-3A_XX | 17 | 2613.9 | 0.029 | Parasite |
| ASV_02714:Bacteriastrum | 13 | 1955.6 | 0.018 | Phototroph |
| ASV_00150:RAD-B-Group-I_X | 12 | 1660.7 | 1.387 | Heterotroph |
| ASV_00897:g__Chlorarachnida_X | 12 | 1074.7 | 0.005 | Mixotroph |
| ASV_00008:Pelagomonas | 11 | 1221.1 | 6.559 | Phototroph |
| ASV_00308:Pedinellales_X | 11 | 1228.3 | 0.016 | Mixotroph |
| ASV_00538:MAST-4A_XX | 11 | 1657.0 | 0.113 | Heterotroph |
| ASV_00632:Minorisa | 11 | 1048.6 | 0.011 | Heterotroph |

### Heterotrophic picoeukaryotes showed stable abundance during the warm season in the ML

After excluding Alveolata, the marine stramenopiles group (MAST) exhibited the highest relative abundance in the ML (Fig. 5B, Fig. 5D). This group of uncultured heterotrophic picoeukaryotes (2–5 µm) consume bacteria and picophytoplankton (Massana et al., 2014; Orsi et al., 2018) and are often highly abundant in the surface layer (Giner et al., 2019; Obiol et al., 2020). It was recently found that MAST require light for metabolic processes associated with heterotrophy due to the presence of rhodopsins (Labarre et al., 2021).

Within MAST, the ASV assigned as MAST-1D showed the highest average abundance (13.3%; Fig. 5B). The ASVs MAST-1D, MAST-1C, MAST-3E, and MAST-4B were positively correlated with temperature (Fig. 6A) and exhibited significantly different abundances (log2fold) in the ML (Fig. 6B), which suggests that they have an affinity for warm and well-lit environments. MAST abundance among all four regions remained stable, although minor changes in abundance were observed in MAST-1C, MAST-D, MAST-3E, and MAST-4B at stations G41 (XIXIMI-4), G44 (XIXIMI-4), and H45 (XIXIMI-4) in the southern region and TS1 (XIXIMI-5) and A10 (XIXIMI-4) in the northern region. Notably, MAST tended to decrease in abundance in stations in which diatom and MOCH-5 abundance increased (Fig. 5D).

**Figure 6.**
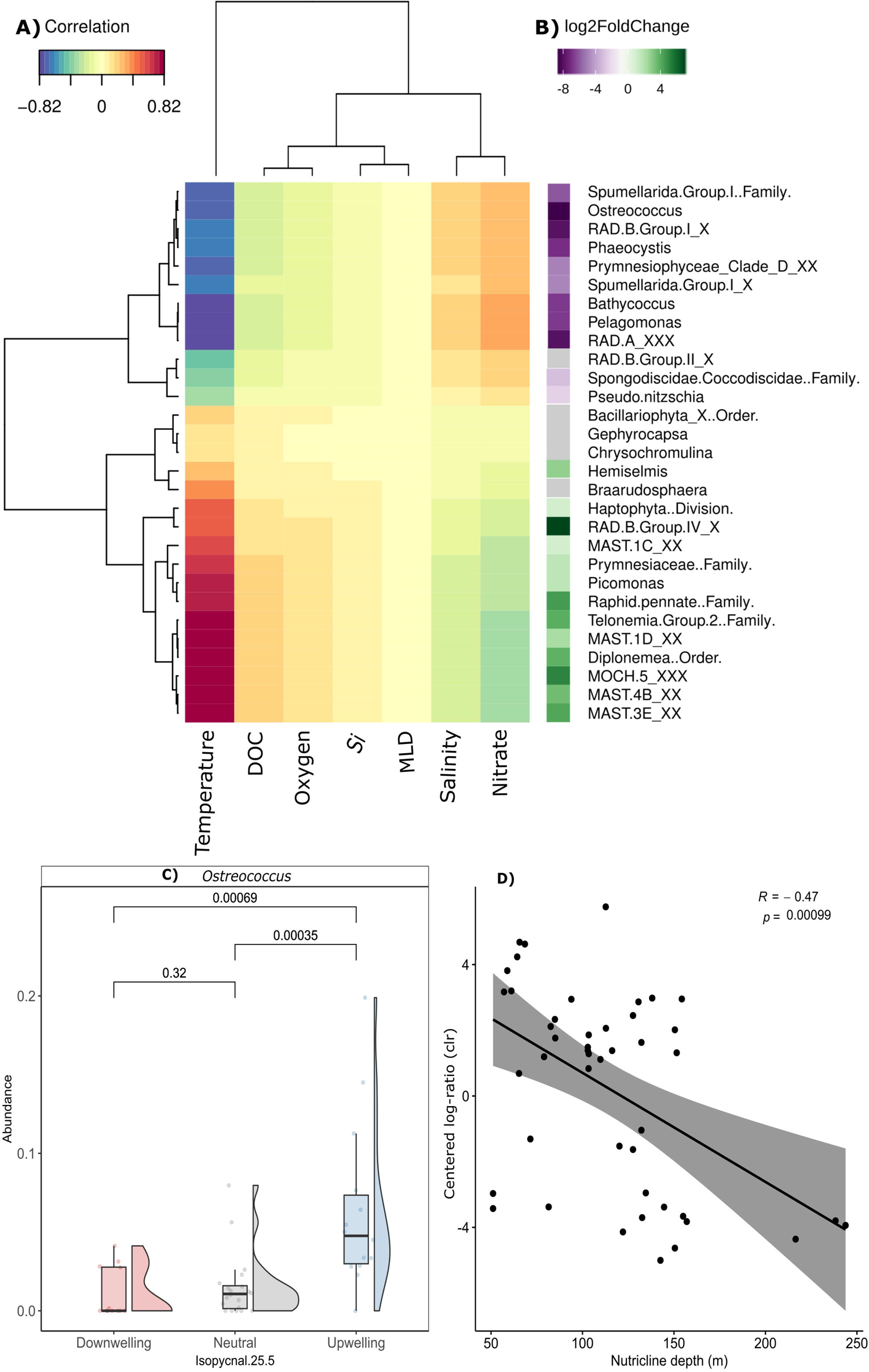
Clustered heatmap representing (A) the correlation between microeukaryote abundance (excluding Alveolata) with environmental parameters. (B) The differential abundance of the principal microeukaryote ASVs) expressed in log2 fold-change with significant differences between the mixed layer d(ML; positive values) and deep chlorophyll maximum (DCM; negative values).. The significance of the ASVs was adjusted (p-value < 0.01). (C) Differences in mean abundance of *Ostreococcus* due to upwelling, downwelling, and neutral conditions, and (D) Spearman correlation of *Ostreococcus* abundance with nutricline depth.

The co-occurrence analysis found that ASV_00425:MAST-3I and ASV_00131:MAST-3E appear to be key species (Table 2). In addition, although MAST-3 was not abundant, it was quite diverse (27 ASVs). This key species is associated with *Solenicola*, which is composed of diatom parasites of the *Leptocylindrus* genus (Gómez et al., 2011) and was recently reported to be associated with diatoms in a bloom event (Ollison et al., 2022). *Solenicola* is also abundant in oligotrophic regions (Lin et al., 2021).

Heterotrophic and photosynthetic bacteria are abundant in the ML (Linacre et al., 2019; Raggi et al., 2020). Previous studies have documented that MAST-4 consumes *Synechoccocus* (Lin et al., 2012) while MAST-9 is strongly correlated with *Acidobacteria* subgruops, *Chloroflexiy* TK10, and *Nitrospinia* (Lin et al., 2021). Also, our data suggest that these MAST may aid in maintaining stable bacterial and picoeukaryote populations at this depth, which may be associated with bacterial grazing (Massana et al., 2014).

### The photosynthetic community exhibits low abundance in the ML

The ASVs affiliated to Haptophyta (Family), Prymnesiphyceae (class), and Prymnesiaceae (Family) are enriched in the ML (Fig. 5C), and different species were present in each depth layer (Fig. 5B, Fig. 5D). Prymnesiophyceae family are diverse and abundant in subtropical oceanic regions (Cortés et al., 2001; Haidar and Thierstein, 2001; Liu et al., 2009). In the Bay of Campeche, *Emiliania huxleyi* and *Gephirocapsa oceanica* have been reported to be equally important diatoms that export carbon to the deep ocean in spring (Hernández-Becerril et al., 2008) and show a well-marked vertical succession of coccolithophores (Prymsesyophyceae order) in the southern region (Baumann and Boeckel, 2013). Furthermore, pigment analyses in the deep-water region of the GoM indicated that Prymnesiophyceae is the most abundant taxa in the euphotic zone after *Prochlorococcus* (Selph et al., 2021).

Coccolithophorids and foraminifera are the main contributors to biogenic calcium carbonate in sediments and input 25–50% of this in the deep-water region of the GoM (Ward, 2017), which makes these taxa central players within the biological pump of the gulf. The ASVs identified as *Gephyrocapsa* were found in abundance in the ML (2.27%) and DCM (2.08%) when compared to those identified as *Braarudosphaera* (3.59%) and *Chrysochromulina* (4.59%), which were found only in the ML (Fig. 5D). These last two taxa are known to live in symbiosis with the diazotrophic cyanobacteria UCYN-A (Endo et al., 2018; Gérikas Ribeiro et al., 2018; Hagino et al., 2013). Moreover, *Chrysochromulina* was found to be a highly diverse genus in the ML (64 ASVs). This mixotroph may shift to bacterial grazing under high irradiance or phosphate-limited conditions (Hansen and Hjorth, 2002; Unrein et al., 2013). In contrast, the non-calcified prymnesiophyte *Phaeocystis,* was significantly more abundant (log2fold) in the DCM (Fig. 6B) than in the ML despite its low abundance in both the DCM and ML (4 and 2% respectively). In addition, its abundance was positively correlated with nitrate and negatively correlated with temperature (Fig. 6A), which indicates that this taxa showed a preference for DCM-like conditions (Sow et al., 2020; Sun et al., 2022).

Our results also show that Haptophytes may play an important role in regulating bacterial and protist populations in the ML, as has been reported for other oligotrophic regions (Endo et al., 2018; Frias-Lopez et al., 2009). Haptophytes contribute to new primary productivity in the DCM and may also participate in photosymbiosis with diazotrophic cyanobacteria in the ML (Krupke et al., 2015), whereas high rates of atmospheric nitrogen (N_2_) fixation have been estimated in the GoM during the summer principally in anticyclonic eddies (Hernández-Sánchez et al., 2021).

Diatom abundance tended to be low in both the ML and DCM. The ASVs identified as Bacillariophyta and the ASVs identified as Raphid-Pennate showed average abundances of 2.6% and 4.5%, respectively, in the ML, and 0.77% and 0.18%, respectively, in the DCM (Fig. 5C, Fig. 5D). Nonetheless, contrary to the typical diatom patterns of high abundance and diversity in cold, nutrient-rich regions (Ramond et al., 2021; Trefault et al., 2021), in the GoM, diatom abundance increased under stratified and oligotrophic conditions in the ML. Bacillariophyta ASVs were more abundant in the southern stations of G41 (39.6%), G44 (26.11%), and H45 (52.9%) during XIXIMI-4. Similarly, the highest ASV abundance values of Raphid-Pennate were identified in stations TS-1 during XIXIMI-5 (44.6%) and A10 during XIXIMI-4 (12.6%; Fig. 5D, Fig. 5E). In contrast, the ASV abundance values of Raphid-Pennate in the DCM in the same stations were less than 1%. However, in station G45 during XIXIMI-4, the ASVs affiliated to Bacillariophyta and Raphid-Pennate showed high abundances (36 and 6% respectively, Fig. 5D). This increase cannot be attributed to mesoscale effects in the ML (Damien et al., 2018b; Lee-Sánchez et al., 2022) given the predominance of oligotrophic conditions and high temperatures. This pattern has been detected in oceanic regions worldwide, with the high diversity of diatom ribotypes (18S-V9) believed to be the result of adaptation to highly stratified conditions (Malviya et al., 2016). In our case, adaptations to ML conditions could be due to their ability to extract nutrients from the nutricline, migrate to shallower depths with greater light levels by regulating buoyancy, and store nitrate within vacuoles, although they may also be due to associations with atmospheric nitrogen-fixing cyanobacteria such as *Rhisozolenia* and *Hemiallus* species associated with *Richellia* (Gómez et al., 2005; Kemp and Villareal, 2013, 2018). The vertical displacement strategy was likely at play in this study because the expected abundance pattern agreed with what was observed in stations in which the nutricline was shallower than 60 m, and some species are capable of migrating that distance in eight hours (Moore and Villareal, 1996).

The reads affiliated to diatoms were mostly assigned to the Bacillariophyta order and Raphid-pennate family, followed by *Chaetoceros* spp. in the ML and *Pseudo-Nizschia*, *Thalassiosira* spp., and Corethron in the DCM, which showed abundance values > 1% (Fig. 5C, Fig. S4). This result agrees with what has been most frequently reported for the GoM based on microscopy analyses (Ghinaglia et al., 2004; Licea et al., 2016; Licea et al., 2011; Merino-Virgilio et al., 2013). The abundance of diatoms of the Raphid-Pennate family was positively correlated with temperature and negatively correlated with nitrate (Fig. 6A). Species of this family exhibited significantly different abundances (log2fold) between the ML and DCM (Fig. 6B). These diatoms are adapted to live in sediments and the water column and can tolerate a wide range of environmental conditions while also exhibiting a high potential for dispersal (Endo et al., 2018; Grippo et al., 2010; Kooistra et al., 2007; Piredda et al., 2018).

We also observed that diatoms of the Raphid-Pennate family are likely being transported by rivers. This was the case for station A10-X4, which was located within a zone influenced by low salinity (33–35) in the northeastern region of the GoM (Fig. S5). Here, pennate diatoms could be transported from the continental shelf to the oceanic region by the Mississippi River plume, which brings low-salinity and nutrient-rich water into the oceanic region of the gulf during summer (Morey et al., 2003; Wawrik and Paul, 2004). On the other hand, station TS-1 exhibited low salinity and could be influenced by fluvial discharge from the Veracruz and Tamaulipas platform (Fig. S5) and by cyclonic circulation (Hamilton et al., 2018; Martínez-López and Zavala-Hidalgo, 2009; Zavala-Hidalgo et al., 2014). Furthermore, the silicate concentrations in the ML were higher in the southern region and station TS1, indicating that both regions are influenced by the coast (Fig. 1, Fig. 2E). In the oceanic region, the silicate concentration was lower than the limiting concentration for diatoms (> 2 µM; Kemp and Villareal, 2018 and references therein). On average, diatom abundance tended to be low in the ML and DCM during summer, although increases in abundance can occur under conditions of high temperature and low nitrate conditions and contribute to high export rates of carbon and silicic acid in the GoM.

### Picoeukaryotes respond to nitrate input in the DCM

In the DCM, the picoeukaryote *Pelagomonas sp.*, which is a member of the Ochrophyta order, was among the most abundant taxa (excluding Alveolata; Fig. 5B, Fig. D) and was represented by only two ASVs (ASV_00008 and ASV_03036) assigned to *P. calceolata*. The abundance of this picoeukaryote was significantly different (log2fold) between the DCM and ML (Fig. 6B) and was positively correlated with nitrate and negatively correlated with temperature (Fig. 6A). *Pelagomonas* has been commonly reported in the DCM in oceanic and oligotrophic regions via pigment, amplicon, and chloroplast genome analyses (Brannock et al., 2016; Selph et al., 2021; Latasa et al., 2017; Wang et al., 2019; Worden et al., 2012) and at the surface under vertically mixed conditions (Choi et al., 2020; Rii et al., 2018). The success of *Pelagomonas* is due to its ability to adapt to conditions of iron limitation (Timmermans et al., 2005) and low irradiance (Dimier et al., 2009; Kang et al., 2021) as well as its small cell size (1–3 µm; Andersen et al., 1993), which allows these picoeukaryotes to quickly uptake nutrients from the environment thanks to their high surface-area-to-volume ratio (Marañón, 2015; Raven, 1998; Ward et al., 2012).

Although iron can be a limiting micronutrient, the DCM layer can be enriched by dissolved Fe and nitrate due to the uplift of isopycnals inside of CE (Hawco et al., 2021). In fertilization experiments, diatom abundance has been found to increase as the concentrations of macro- and micronutrients increase (Alexander et al., 2015; Fawcett et al., 2011). In the Gulf of Mexico, the co-limitation of macro- and micronutrients could inhibit diatom proliferation in the DCM, which would favor picoeukaryote species like *Pelagomonas*. This has been reported at the edges of oceanic gyres in the south Atlantic and subtropical Pacific, where co-limitation has been found to be conducive to a high diversity and abundance of picoeukaryotes such as Pelagophytes (Browning et al., 2017, 2022). Furthermore, studies using metatranscriptomics approaches have observed that *Pelagomonas* are associated with nitrate assimilation in the DCM (Dupont et al., 2015) and that they may regulate the relative expression levels of ferrodoxin and flavodoxin depending on the iron concentration (Carradec et al., 2018).

Another interesting result was the presence of the green alga (Chlorophyta) *Ostreocuccus sp.*, which was represented by 2 ASVs (ASV_00123 and ASV_02103). The average abundance was low (3.16%), although this increased in stations under the influence of upwelling conditions with relatively shallow nutriclines such as station A10-X4 (Fig. 5D). This station was located inside a CE between the LC and an LCE and exhibited a nutricline depth of 59 m. The abundance of *Ostreococcus* sp. in this station reached 27.8%. Surprisingly, *Ostreocococcus* was the only genus that showed significant differences in mean abundance between upwelling and downwelling (p = 0.00084) and upwelling and neutral (p = 0.0003) conditions in the DCM (Fig. 6C). Also, the abundance of this genus was significantly different (log2fold) in the DCM compared to that of the ML (Fig. 6B).

Like *Pelagomonas*, *Ostreocococcus* was positively correlated with nitrate and negatively correlated with temperature (Fig. 6A). *Ostreococcus* has been commonly reported to be abundant in nutrient-rich areas such as those in coastal regions, and constrained in the DCM in oligotrophic regions (Tragin and Vaulot, 2018; Worden et al., 2006). Although, it is dominant in subpolar boundary regions in the North and South Pacific and North Atlantic (Bolaños et al., 2020; Clayton et al., 2017; Gutierrez-Rodriıguez et al., 2021). In addition, this genus is also abundant in the Gulf Stream (Demir-Hilton et al., 2011) and in the San Pedro Channel in the Pacific into the DCM (Countway and Caron, 2006).

*Ostreococcus*, like *Prochlorococcus,* shows niche partitioning with ecotypes that are adapted to high and low light conditions (Johnson et al., 2006; Moore and Chisholm, 1999). The *Ostreococcus* OI ecotype is associated with mesotrophic and coastal conditions, and the *Ostreocococcus* OII ecotype is abundant in the DCM of oligotrophic regions (Demir-Hilton et al., 2011; Simmons et al., 2016). In this study, the V9 region of the 18S rRNA gene did not provide sufficient resolution to distinguish ecotypes (Monier et al., 2016); however, the results clearly show that *Ostreococcus* increased in abundance in the DCM when the nutricline upwelled (Rho = −0.47, p = 0.00099, Fig. 6D).

Previous studies in the oceanic region of the GoM have reported that primary productivity tends to increase in the DCM during the warm season (Pasqueron de Fommervault et al., 2017; Yingling et al., 2021). Our data suggest that the main microbial eukaryotic genera contributing to new primary production in the DCM during the warm season were principally picoeukaryotes, such as *Pelagomonas*, *Ostreococcus*, and *Bathiococcus*, and the haptophyte *Phaeocystis.* These groups contribute to the primary productivity of other low light ecotypes of *Prochlorocococcus* (Linacre et al., 2019). In the oligotrophic regions of the Pacific and Atlantic, photosynthetic picoeukaryotes in the DCM have been reported to uptake upwelled nitrate as their primary nitrogen source (Fawcett et al., 2011; Painter et al., 2014; Rii et al., 2018).

In the deep-water region of the GoM (> 1000 m), cytometry analysis has revealed an increase in the biomass of picoeukaryotes in cyclonic mesoscale structures during winter (Linacre et al., 2015). It has been reported that within CEs in the subtropical Pacific, the uplift of isopycnals reduces light stress while increasing the nitrate and dissolved Fe concentrations, which in turn generates favorable conditions for the proliferation of picoeukaryotes during the mature phase of the CE (Hawco et al., 2021). In this study, the major difference in community structure between upwelling and downwelling stations in the DCM was determined by the increased abundance of *Ostreococcus sp.* when the nutricline was shallow (Fig 6C, Fig. 6D). Therefore, *Ostreococcus* may be an indicator of elevations in the nutricline during summer because nitrate can be immediately removed by picoplankton.

During summer in the four XIXIMI cruises included in this study, we observed that nutrients did not penetrate the ML due to stratification (Lee-Sánchez et al., 2022), while the microbial loop was dominant (Selph et al., 2021; Linacre et al., 2019), and the effects of mesoscale structures were observed in the DCM (Pasqueron de Fommervault et al., 2017; Lee-Sánchez et al., 2022). Unlike what was recorded in the subtropical Pacific and Atlantic during the initial CE intensification phase with well-defined mixed layers and large diatom blooms in the DCM due to macro and micronutrient enrichment (Brown et al., 2008; McGillicuddy et al., 2007), the abundance of diatoms was very low, and photosynthetic picoeukaryotes appeared to respond to upwelled nitrate. In fact, in the subtropical Pacific and Mediterranean, communities in the DCM similar to those found in this study composed mainly of Haptophytas and photosynthetic picoeukaryotes have been reported in mature CEs (Belkin et al., 2022; Hawco et al., 2021).

The success of picoeukaryotes over diatoms in the DCM could be due to the vertical diffusive flux of nitrate, which has been previously estimated to be very low in the deep-water region of the GoM during summer (Damien et al., 2021; Kelly et al., 2021). This process supplies new nitrate to phytoplankton (Villamaña et al., 2019) and could confer an advantage to picophytoplankton over diatoms with regard to the uptake of nutrients in low concentrations from the environment (Damien et al., 2021; Marañón et al., 2001; Villamaña et al., 2019) when the N:P ratio is low and ammonium is the predominant source of nitrogen (Rodríguez-Gómez et al., 2022; Yingling et al., 2021).

Silicate is another limiting factor for diatom blooms in the central, northern, and LC regions of the GoM (Fig. 2) when its concentration is lower than 2 µM (Kemp and Villareal, 2018). Our data showed this low diatom abundance pattern in the region where Si was limited. The increase in the deep penetration of photosynthetically active radiation in the subsurface layer in highly stratified environments (Agusti et al., 2019), which increases the abundance of photosynthetic picoeukaryotes in subtropical regions and increases the probability of a deep picoeukaryote bloom as temperature rises, could also favor picoeukaryotes over diatoms, which typically prefer turbulent environments (Cullen et al., 2007). Given these results, some questions arise. Is macro and micronutrient (e.g., nitrate-iron) co-limitation present in the DCM? Why was diatom dominance not observed when the nutricline was shallow in the DCM during the warm season?

### Non-constitutive mixotrophs are abundant in the DCM

Another group that was highly abundant after Alveolata in the DCM was siliceous Rhizaria. An overestimation of siliceous Rhizaria abundance is expected with metabarcoding analysis because this taxa, like Alveolata, has a high rDNA gene copy number (Biard et al., 2017; de Vargas et al., 2015), although large cell volumes positively correlate with read abundance (Biard et al., 2016; Llopis Monferrer et al., 2022). This group is associated with high carbon and biogenic silica export (Guidi et al., 2016), and its abundance tends to increase in mesopelagic regions (Countway et al., 2007; Georges et al., 2014). The Spumellaria-Group-I (Family) was represented by 11 ASVs and was the most abundant (11%; Fig. 5B, Fig. 5D). The abundance of this taxa was significantly different in the DCM when compared to that of the ML (Fig. 6B). It has also been reported to be highly dominant in sediment traps (Gutierrez-Rodriguez et al., 2019). Radiolaria are non-constitutive mixotrophs (Flynn et al., 2019) associated with different photosynthetic species such as Chlorophytes, Haptophytes, and Prasinophyteas (Biard, 2022 and references therein). Interestingly, the co-occurrence analysis showed only a negative association of Spumellarida-Group-I with Bacillariophyta (ASV_00786; Supporting information, Table S4), despite being the most abundant Rhizaria in the DCM.

The environmental clades RAD-A and RAD-B (radiolarian groups, Taxopodida order) exhibited opposing distribution patterns between the ML and DCM, where RAD-B-group-IV (4 ASVs) showed the highest abundance in the ML compared to that of the DCM, and RAD-B-group-I (9 ASVs) and RAD-A (21 ASVs) showed the highest abundance in the DCM compared to that of the ML (Fig. 6B). The Taxopodia order does not exhibit symbiotic associations and very little is known about its ecology (Biard, 2022). RAD-A has been recorded with high metabolic activity in the DCM, and RAD-B has been found to increase its activity with depth (Giner et al., 2019). RAD-B is found in the mesopelagic region (Gutiérrez-Rodríguez et al., 2022) and is associated with particles (Duret et al., 2020). Our co-occurrence analysis showed positive associations between *RAD-B Group II* with RAD-B Group IV and RAD-A with RAD-B Group-IV (Fig. 7c) and negative associations between RAD-A with Raphid-pennate and RAD-A with Bacillaryophyta_X, which could indicate the presence of amensalism. In this case, RAD-A probably consumes the DOM produced by phytoplankton, but it remains unclear how diatoms could be affected. In addition, a negative relationship between RAD-B Group-IV and Syndiniales was found that suggested parasitism (Fig. 7A, Fig. 7B). Nonetheless, these associations need to be validated with other techniques such as microscopy.

**Figure 7.**
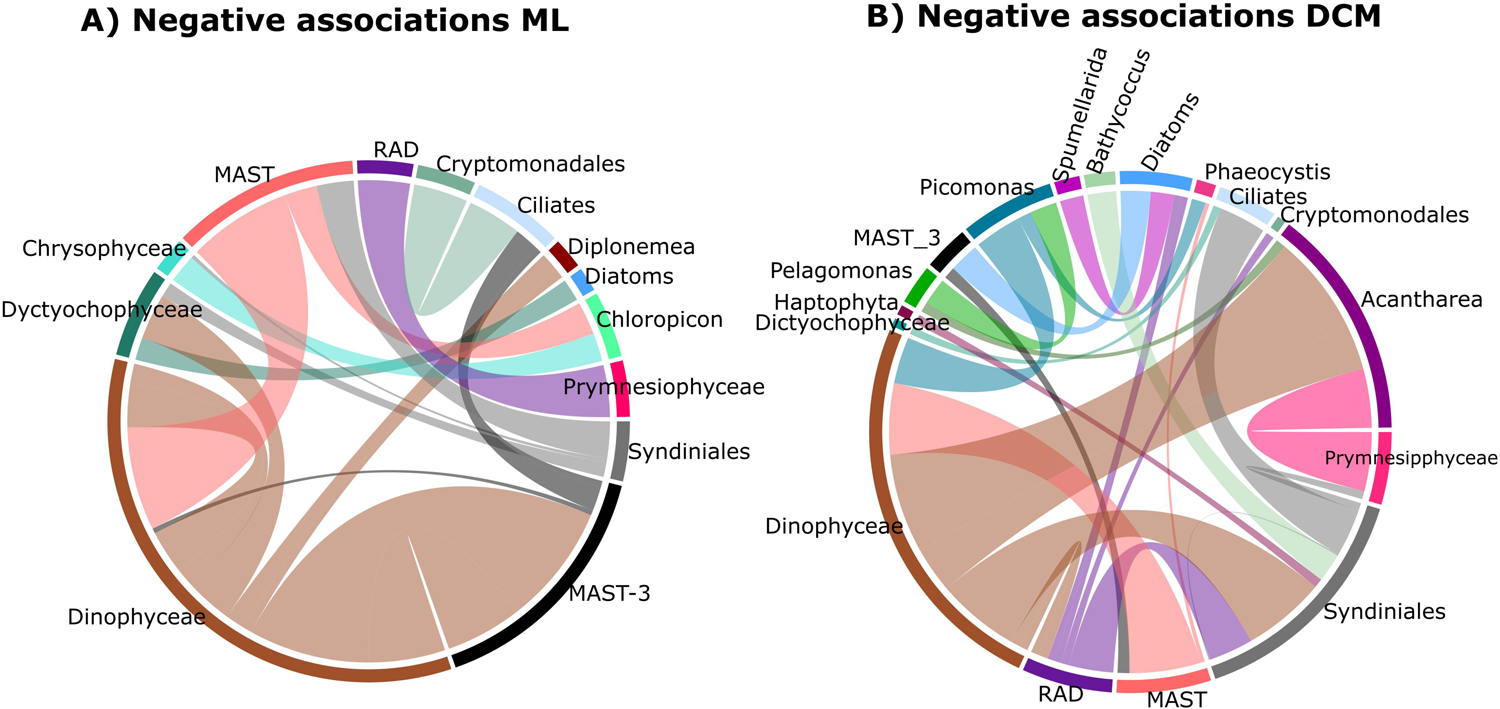
Protist ASVs interaction network Negative associations in (A) the mixed layer (ML) and (B) the deep chlorophyll maximum (DCM). Ribbon Ribbons represent the proportions of nodes (ASVs).

### Top-down control strongly influences the protist community of the euphotic zone

Our results suggest that abiotic factors, such as temperature and nutrient concentrations, have smaller effects on the protist community than do biological factors, such as top-down controls, which play major roles in shaping the community structure in both the ML and DCM. Protist-protist co-presence (positive ecological association) and self-exclusion (negative ecological association) interactions have been found to increase with temperature in oligotrophic regions in which DOM is rapidly recycled (Armengol et al., 2019; Azam et al., 1983). We found that positive interactions were 70% (571) in the ML and 62% (514) in the DCM. This increase in positive protist-protist interactions in the ML could be due to the establishment of symbiotic or co-existence relationships in which protists share the same niche.

The co-occurrence network revealed high numbers of positive ecological associations among the Dinophycea class with other dinoflagellates, diatoms, and ciliates in the ML and DCM (e.g., *Gymnodinium*-*Torodinium* and Gymnodiaceae-Ciliophora; Fig. S6). However, the positive associations of predator and prey (Karlodinium-Picomonas) and parasite and host (Syndiniales-Hemiselmis) suggest that the co-presence of these relationships might not affect mortality (Faust and Raes, 2012; Fuhrman et al., 2015). This co-presence may be associated with a stable community assembled over time (i.e., coevolution; Mougi and Iwasa, 2011), with reciprocal selective pressure and rapid adaptation (Liu et al., 2022), which would indicate positive relationships between resident species (e.g., Gymnodiniales) that exhibit high abundance year-round (> 40%: Linacre, et al., 2021).

Self-exclusion was high in both the ML (30%, 246) and DCM (38%, 315) when compared to what has been found in other studies, which have reported self-exclusion values lower than 10%, although it is important to note that the results and data were obtained with different methods in other studies (Krabberød et al., 2021; Lima-Mendez et al., 2015; Liu et al., 2022). Under oligotrophic conditions, a high number of protist-protist mutual exclusion associations would be expected, such as parasitism, competition, and predation, in ecosystems in which the microbial loop dominates (Azam and Graf, 1983).

In the ML, the predominant negative interaction was parasitism: MAST-3 parasitizing dinoflagellates and ciliates (Fig. 7A). In the DCM, in addition to parasitism by Syndinials with negative associations with dinoflagellates, ciliates, haptophytas, and *Bathyococcus* (Fig. 7B), we also found the co-occurrence of possible predation (e.g., Pelagomonas-Picomonas), resource-based competition (e.g., Gyrodinium and Karlodinium), and even parasitism-parasitism such as with Dino-Group-II-Clade-4-Dino-Group-II-Clase 26 (Syndinial-Syndinial). Finally, the keystone species were predominantly mixotrophic and heterotrophic protists at both depths (Table 2). The associated ASVs exhibited a high degree of betweenness but low abundance (Table 2), indicating that hub and connector species were present that are highly connected to other protist species in the ecosystem (Krabberød et al., 2022). This suggests that protists are key components of the GoM ecosystem that perform important functions while influencing the food web and biogeochemical cycles.

## Conclusions

We explored the composition of the protist community during the warm season in the deep-water region of the GoM. For the first time, metabarcoding analysis allowed us to broadly explore the community of the GoM MEEZ to improve our understanding of the fraction of the community that cannot be detected by conventional methods. Our results showed that the ML is dominated by mixotrophic and heterotrophic picoeukaryotes while the autotrophic community is poorly represented. In contrast, the DCM exhibited a community dominated by photosynthetic picoeukaryotes (after excluding alveolates). Surprisingly, *Ostroecoccus* was the unique genus that tended to increase in abundance when the nutricline was shallow. However, top-down controls, such as parasitism, competition, and predation, may exert a strong influence over the abundance of protists in the ML and DCM. We recommend exploring the functions of the protist community using a metatranscriptomics approach to better understand how this community responds to the supply of macro and micronutrients and the functions that dominate the community at both depths. A network analysis that includes the prokaryotic fraction would also provide valuable information of the energy flow of the whole microbial community. This study provides an essential baseline of the composition of the protist community in the southern GoM and demonstrates how this community responds to environmental and biological factors in a highly oligotrophic and warm ecosystem.

## Supporting information

Table S4. Environmental details of the samples

Table S5. ASVs co-occurrence associations from the ML

Table S6. ASVs co-occurrence associations from the DCM

Table S7. ASV count table with taxonomy from the 35 top taxa.

Supplementary figures and statistic tabless.

## Acknowledgements

We thank the scientific participants and crew of R*/V Justo Sierra* (UNAM) for their assistance during the oceanographic campaigns. We are grateful to Uriel Mirabal for providing data of the mixed layer depth. Finally, we acknowledge PEMEX’s specific request to the Hydrocarbon Fund to address the environmental effects of oil spills in the Gulf of Mexico.

## Declarations

N/A

## Author contributions

Conceptualization: A L-L, K S-C; Formal analysis: K S-C, MA M-M; Funding acquisition: A L-L; Investigation: K S-C, J C-R, Y O-S; Methodology: A L-L, K S-C, J C-R; Project administration: A L-L; Resources: A L-L, V C-I; Supervision: A L-L, L L, V C-I; Validation: A L-L;; Writing – original draft: K. S-C; Writing – review & editing: MA M-M, A L-L, J C-R, Y O-S, L L, V C-I

## Data availability

The raw sequence data that support these findings were deposited in the NCBI database under accession number PRJNA971952.

## Funding

This research was supported by the Mexican National Council for Science and Technology (CONACyT)-Mexican Ministry of Energy (SENER)-Hydrocarbon Fund (project 201441). CONACYT awarded K. S-C with a Ph.D. scholarship (474503). This is a contribution of the Gulf of Mexico Research Consortium (CIGoM).

## Conflict of interest

The authors declare no conflict of interest. The funders had no role in the design of the study; in the collection, analyses, or interpretation of data; in the writing of the manuscript; or in the decision to publish the results.

## Ethics approval, Patient consent, Permission to reproduce material from other sources, and Clinical trial registration

N/A

## Supporting information

**Table S1.** Environmental details of the samples. Pressure. Temperature. Salinity. Sigma-t. Oxygen. Fluorescence. Latitude. Longitude. MLD. Nutricline. Isopycnal 25.5. Nitrate. Si. DOC.

**Table S2.** ASVs co-occurrence associations from the MLD

**Table S3**. ASVs co-occurrence associations from the DCM

**Table S4**. ASV count table with taxonomy.

**Appendix S1**. Supplementary figures and statistic tables.

## Notes

### Competing Interest Statement

The authors have declared no competing interest.

