## Supplementary material for "Response of microbial eukaryote community to the oligotrophic waters of the Gulf of Mexico: a plausible scenario for warm and stratified oceans": Table S4. Environmental details of the samples

| Station | Depth | Cruise | Region | Pressure (db) | Tem. (°C) | Abs. Salinity (g/kg) | sigma.t ($kg.m^3$) | Oxigen (µmol.kg) | Fluor. (RFU) | Latitude (°N) | Longitude (°W) | MLD (m) | Nutri. (m) | Iso. 25.5 (m) | Nitrito (µmol. $Kg^{-1}$) | Si (µmol. $Kg^{-1}$) | DOC (µmol. $Kg^{-1}$) |
|---|---|---|---|---|---|---|---|---|---|---|---|---|---|---|---|---|---|
| A10_X4 | DCM | XIXIMI-4 | North | 33.60 | 26.35 | 36.47 | 24.03 | 193.98 | 1.37 | 25.01231 | -86.99612 | 33.80 | 58.90 | Upwelling | | | |
| A3_X4 | DCM | XIXIMI-4 | North | 100.00 | 24.23 | 36.42 | 24.65 | 219.59 | 1.44 | 25.00487 | -93.99754 | 45.70 | 134.60 | Donwelling | 1.45 | 1.29 | 63.60 |
| A5_X4 | DCM | XIXIMI-4 | North | 104.80 | 23.14 | 36.53 | 25.05 | 178.09 | 1.78 | 25.00098 | -91.99740 | 46.50 | 122.06 | Donwelling | 1.19 | 1.15 | 58.50 |
| A7_X4 | DCM | XIXIMI-4 | North | 70.60 | 21.42 | 36.42 | 25.46 | 198.77 | 1.87 | 24.99644 | -90.00124 | 18.90 | 68.51 | Upwelling | 2.13 | 1.74 | 61.10 |
| B14_X4 | DCM | XIXIMI-4 | North | 108.60 | 25.69 | 36.56 | 24.31 | 181.30 | 1.62 | 24.00223 | -92.30044 | 24.80 | 132.20 | Donwelling | 0.37 | 1.07 | 64.70 |
| B16_X4 | DCM | XIXIMI-4 | North | 80.90 | 23.44 | 36.43 | 24.89 | 204.19 | 1.68 | 24.00002 | -90.00066 | 35.80 | 110.20 | Donwelling | 0.30 | 1.47 | 65.70 |
| B17_X4 | DCM | XIXIMI-4 | North | 78.70 | 21.92 | 36.41 | 25.32 | 212.78 | 1.78 | 24.00030 | -89.00124 | 24.80 | 79.00 | Neutral | 0.58 | 1.49 | 67.40 |
| F39_X4 | DCM | XIXIMI-4 | Center | 74.40 | 20.59 | 36.45 | 25.71 | 167.18 | 1.90 | 21.00273 | -93.00302 | 27.70 | 61.13 | Upwelling | 5.25 | 0.96 | 54.20 |
| G41_X4 | DCM | XIXIMI-4 | South | 78.00 | 21.21 | 36.59 | 25.65 | 161.68 | 1.80 | 20.49662 | -95.50254 | 22.90 | 65.30 | Upwelling | | | |
| G44_X4 | DCM | XIXIMI-4 | South | 37.00 | 27.64 | 36.58 | 23.70 | 195.60 | 1.91 | 20.50070 | -92.59590 | 26.70 | 53.60 | Upwelling | 0.09 | 0.48 | 77.50 |
| H45_X4 | DCM | XIXIMI-4 | South | 24.40 | 29.91 | 36.46 | 22.86 | 193.37 | 1.69 | 19.99906 | -95.60076 | 22.80 | 51.05 | Upwelling | | | |
| Y1_X4 | DCM | XIXIMI-4 | LCs | 54.40 | 27.75 | 36.49 | 23.60 | 190.79 | 1.42 | 21.66822 | -86.37366 | 47.60 | 71.50 | Neutral | | | |
| Y4_X4 | DCM | XIXIMI-4 | LCs | 79.00 | 27.30 | 36.41 | 23.69 | 196.40 | 1.40 | 21.75890 | -85.93483 | 28.80 | 144.55 | Donwelling | 0.29 | 1.05 | 68.30 |
| A10_X5 | DCM | XIXIMI-5 | LCs | 95.60 | 25.70 | 36.29 | 24.10 | 208.23 | 0.65 | 24.86036 | -87.07192 | 25.90 | 166.85 | Donwelling | 0.20 | 1.56 | 75.82 |
| A4_X5 | DCM | XIXIMI-5 | North | 90.20 | 22.61 | 36.39 | 25.10 | 216.20 | 0.73 | 25.01360 | -93.01217 | 30.80 | 112.74 | Neutral | 0.60 | 1.40 | 64.49 |
| A7_X5 | DCM | XIXIMI-5 | North | 96.20 | 25.00 | 36.51 | 24.48 | 201.86 | 0.66 | 25.02784 | -90.05195 | 21.80 | 127.63 | Neutral | | | |
| B12_X5 | DCM | XIXIMI-5 | North | 80.20 | 23.64 | 36.43 | 24.83 | 224.57 | 0.28 | 23.99479 | -95.01317 | 20.90 | 150.48 | Donwelling | 0.19 | 1.75 | 76.18 |
| B17_X5 | DCM | XIXIMI-5 | North | 89.90 | 25.56 | 36.48 | 24.29 | 207.19 | 0.67 | 24.01060 | -89.03952 | 18.90 | 109.77 | Neutral | 0.03 | 1.10 | 70.05 |
| B18_X5 | DCM | XIXIMI-5 | North | 74.20 | 20.67 | 36.51 | 25.73 | 201.01 | 0.75 | 23.98566 | -86.71884 | 22.85 | 57.13 | Upwelling | 2.82 | 1.95 | 65.06 |
| C22_X5 | DCM | XIXIMI-5 | Center | 112.10 | 22.20 | 36.38 | 25.21 | 212.68 | 0.55 | 22.98534 | -94.54785 | 23.90 | 138.08 | Neutral | 0.88 | 1.47 | 60.70 |
| G44_X5 | DCM | XIXIMI-5 | South | 79.90 | 21.58 | 36.44 | 25.43 | 193.43 | 0.90 | 20.53472 | -92.51313 | 18.88 | 81.48 | Upwelling | 0.08 | 3.42 | 86.07 |
| H47_X5 | DCM | XIXIMI-5 | South | 78.60 | 23.02 | 36.38 | 24.97 | 213.74 | 0.50 | 20.03050 | -94.02260 | 18.90 | 103.34 | Neutral | 0.10 | 1.59 | 77.02 |
| PO1_X5 | DCM | XIXIMI-5 | LCs | 118.40 | 25.72 | 36.27 | 24.08 | 208.97 | 0.69 | 25.47465 | -88.02782 | 26.80 | 254.68 | Donwelling | 0.03 | 1.44 | 65.67 |
| TS1_X5 | DCM | XIXIMI-5 | North | 72.60 | 20.98 | 36.43 | 25.59 | 205.13 | 0.56 | 25.75303 | -95.54745 | 18.90 | 65.56 | Upwelling | 0.58 | 1.51 | 67.34 |
| A10_X6 | DCM | XIXIMI-6 | LCs | 114.60 | 27.03 | 36.43 | 23.78 | 192.20 | 0.40 | 24.94308 | -87.05574 | 41.50 | 216.53 | Donwelling | 0.00 | 1.28 | 62.52 |
| B17_X6 | DCM | XIXIMI-6 | North | 96.20 | 22.37 | 36.43 | 25.20 | 163.20 | 0.49 | 24.00114 | -89.00090 | 27.70 | 102.87 | Neutral | 1.73 | 2.45 | 74.48 |
| C21_X6 | DCM | XIXIMI-6 | Center | 99.90 | 22.66 | 36.38 | 25.08 | 181.70 | 0.65 | 22.99944 | -95.50146 | 46.50 | 112.76 | Neutral | 0.39 | 1.72 | 94.69 |
| C25_X6 | DCM | XIXIMI-6 | Center | 110.70 | 22.09 | 36.45 | 25.30 | 176.87 | 0.40 | 22.99957 | -91.00437 | 27.70 | 116.20 | Neutral | 0.00 | 1.68 | 80.63 |
| D27_X6 | DCM | XIXIMI-6 | Center | 90.60 | 23.63 | 36.46 | 24.86 | 177.85 | 0.50 | 22.00038 | -96.00232 | 27.70 | 103.38 | Neutral | 0.40 | 1.82 | 85.02 |
| E33_X6 | DCM | XIXIMI-6 | Center | 100.50 | 22.10 | 36.38 | 25.24 | 189.09 | 0.45 | 21.49064 | -94.50118 | 25.70 | 102.86 | Neutral | 1.71 | 1.87 | 75.72 |
| F37_X6 | DCM | XIXIMI-6 | Center | 88.30 | 22.84 | 36.40 | 25.04 | 180.34 | 0.68 | 21.00914 | -94.99818 | 28.70 | 103.38 | Neutral | 0.00 | 1.65 | 85.83 |
| G40_X6 | DCM | XIXIMI-6 | South | 79.00 | 22.68 | 36.41 | 25.10 | 157.59 | 0.71 | 20.50124 | -96.00020 | 20.80 | 85.10 | Upwelling | 1.81 | 2.12 | 74.71 |
| G44_X6 | DCM | XIXIMI-6 | South | 65.30 | 21.75 | 36.53 | 25.45 | 130.00 | 1.17 | 20.52278 | -92.50004 | 22.80 | 64.30 | Upwelling | 2.64 | 2.06 | 87.65 |
| H45_X6 | DCM | XIXIMI-6 | South | 86.40 | 21.69 | 36.51 | 25.45 | 134.66 | 0.53 | 19.99902 | -95.60639 | 18.80 | 82.60 | Upwelling | 2.57 | 2.47 | 86.65 |
| Y3_X6 | DCM | XIXIMI-6 | LCs | 65.30 | 27.24 | 36.62 | 23.86 | 186.30 | 0.48 | 21.64423 | -86.23118 | 18.80 | 120.17 | Neutral | 0.16 | 1.20 | 78.14 |
| Y7_X6 | DCM | XIXIMI-6 | LCs | 90.30 | 27.30 | 36.49 | 23.75 | 190.53 | 0.54 | 21.66242 | -85.94704 | 33.60 | 154.28 | Donwelling | 0.00 | 1.25 | 80.73 |
| A10_X7 | DCM | XIXIMI-7 | LCs | 101.90 | 26.80 | 36.31 | 23.77 | 192.73 | 0.22 | 24.99736 | -86.99794 | 52.65 | 238.32 | Donwelling | 0.10 | 1.36 | 86.27 |
| A2_X7 | DCM | XIXIMI-7 | North | 99.30 | 23.27 | 36.53 | 25.01 | 203.74 | 0.27 | 24.87534 | -94.97718 | 25.83 | 151.47 | Neutral | 0.24 | 1.64 | 59.61 |
| A6_X7 | DCM | XIXIMI-7 | North | 73.90 | 21.81 | 36.42 | 25.35 | 206.27 | 0.33 | 25.00125 | -91.00100 | 30.80 | 84.93 | Upwelling | 0.10 | 1.27 | 70.58 |
| A7_X7 | DCM | XIXIMI-7 | North | 84.50 | 26.82 | 36.18 | 23.67 | 195.67 | 0.33 | 21.68545 | -85.93707 | 20.86 | 93.87 | Upwelling | 0.97 | 1.63 | 66.84 |
| B12_X7 | DCM | XIXIMI-7 | North | 98.70 | 23.58 | 36.53 | 24.93 | 205.78 | 0.39 | 23.99973 | -95.06710 | 26.83 | 155.00 | Neutral | 0.05 | 1.47 | 70.04 |
| C24_X7 | DCM | XIXIMI-7 | Center | 107.10 | 22.68 | 36.48 | 25.15 | 202.56 | 0.16 | 23.00268 | -91.99990 | 28.82 | 130.63 | Neutral | 0.29 | 1.64 | 78.79 |
| Y6_X7 | DCM | XIXIMI-7 | LCs | 87.60 | 27.20 | 36.32 | 23.65 | 194.83 | 0.15 | 21.65724 | -86.06038 | 93.40 | 142.56 | Neutral | | | |

**Table S4. Environmental details of the samples (continued).**

| Station | Depth | Cruise | Region | Pressure (db) | Tem. (°C) | Abs. Salinity (g/kg) | sigma.t (kg.m3) | Oxigen (µmol.kg) | Fluor. (RFU) | Latitude (°N) | Longitude (°W) | MLD (m) | Nutri. (m) | Iso. 25.5 (m) | Nitrito (µmol. Kg-1) | Si (µmol. Kg-1) | DOC (µmol. Kg-1) |
|---|---|---|---|---|---|---|---|---|---|---|---|---|---|---|---|---|---|
| Y7_X7 | DCM | XIXIMI-7 | LCs | 84.50 | 26.82 | 36.18 | 23.67 | 195.67 | 0.24 | 21.68545 | -85.93707 | 36.70 | 156.96 | Neutral | | | |
| Y9_X7 | DCM | XIXIMI-7 | LCs | 126.10 | 26.72 | 36.24 | 23.75 | 193.82 | 0.18 | 20.75627 | -85.61984 | 45.71 | 243.85 | Donwelling | 0.00 | 1.50 | 81.20 |
| A10_X4 | ML | XIXIMI-4 | North | 6.00 | 29.69 | 36.07 | 22.64 | 194.27 | 1.18 | 25.00738 | -86.99717 | 33.80 | 58.90 | Upwelling | 0.02 | 0.74 | 75.90 |
| A5_X4 | ML | XIXIMI-4 | North | 5.00 | 30.28 | 36.58 | 22.82 | 193.09 | 1.23 | 25.00070 | -91.99882 | 46.50 | 122.06 | Donwelling | 0.06 | 1.00 | 68.20 |
| A7_X4 | ML | XIXIMI-4 | North | 6.00 | 30.03 | 35.65 | 22.20 | 193.48 | 1.44 | 24.99897 | -90.00068 | 18.90 | 68.51 | Upwelling | 0.33 | 1.62 | 78.70 |
| B14_X4 | ML | XIXIMI-4 | North | 6.00 | 30.12 | 36.41 | 22.75 | 194.13 | 1.13 | 24.00060 | -92.30047 | 24.80 | 132.20 | Donwelling | 0.00 | 0.99 | 70.80 |
| B17_X4 | ML | XIXIMI-4 | North | 5.00 | 30.22 | 36.35 | 22.66 | 192.74 | 1.18 | 23.99938 | -89.00152 | 24.80 | 79.00 | Neutral | 0.04 | 1.14 | 78.30 |
| G41_X4 | ML | XIXIMI-4 | South | 6.00 | 29.83 | 36.57 | 22.97 | 194.72 | 1.25 | 20.49970 | -95.49983 | 22.90 | 65.30 | Upwelling | | | |
| G44_X4 | ML | XIXIMI-4 | South | 3.00 | 29.36 | 36.57 | 23.13 | 208.28 | 1.87 | 20.50253 | -92.59746 | 26.70 | 53.60 | Upwelling | 0.12 | 0.53 | 86.40 |
| H45_X4 | ML | XIXIMI-4 | South | 6.00 | 29.71 | 36.18 | 22.71 | 195.01 | 1.78 | 21.50072 | -96.50000 | 22.80 | 51.05 | Upwelling | | | |
| Y4_X4 | ML | XIXIMI-4 | LCs | 7.00 | 30.40 | 36.32 | 22.58 | 196.44 | 1.03 | 21.75537 | -85.93380 | 28.80 | 144.55 | Donwelling | 0.06 | 1.57 | 72.90 |
| A10_X5 | ML | XIXIMI-5 | LCs | 4.00 | 29.48 | 36.45 | 22.99 | 208.66 | 0.10 | 24.88438 | -87.03750 | 25.90 | 166.85 | Donwelling | 0.00 | 1.46 | 74.60 |
| A4_X5 | ML | XIXIMI-5 | North | 9.00 | 29.67 | 35.94 | 22.54 | 203.78 | 1.16 | 25.01125 | -93.00463 | 30.80 | 112.74 | Neutral | 0.01 | 1.31 | 73.90 |
| A7_X5 | ML | XIXIMI-5 | North | 4.00 | 29.10 | 36.44 | 23.12 | 211.19 | 0.99 | 25.00525 | -90.02828 | 21.80 | 127.63 | Neutral | 0.02 | 1.12 | 65.51 |
| B12_X5 | ML | XIXIMI-5 | North | 5.00 | 29.25 | 36.53 | 23.13 | 207.36 | 1.15 | 23.99947 | -95.00952 | 20.90 | 150.48 | Donwelling | 0.19 | 1.75 | 76.18 |
| B17_X5 | ML | XIXIMI-5 | North | 4.00 | 28.99 | 36.34 | 23.07 | 208.66 | 0.88 | 24.00882 | -89.02607 | 18.90 | 109.77 | Neutral | 0.18 | 1.11 | 64.10 |
| B18_X5 | ML | XIXIMI-5 | North | 4.00 | 29.51 | 36.37 | 22.93 | 208.42 | 0.52 | 23.99147 | -86.71227 | 22.85 | 57.13 | Upwelling | 0.15 | 0.96 | 67.47 |
| G44_X5 | ML | XIXIMI-5 | South | 4.00 | 29.52 | 36.00 | 22.64 | 210.43 | 0.58 | 20.52488 | -92.51078 | 18.88 | 81.48 | Upwelling | 2.43 | 1.88 | 66.83 |
| H46_X5 | ML | XIXIMI-5 | South | 4.00 | 29.36 | 35.80 | 22.55 | 210.10 | 1.10 | 19.99460 | -95.00808 | 18.90 | 71.55 | Upwelling | 0.04 | 1.57 | 88.38 |
| H47_X5 | ML | XIXIMI-5 | South | 6.00 | 28.85 | 36.05 | 22.91 | 210.95 | 0.38 | 20.01670 | -94.01348 | 18.90 | 103.34 | Neutral | 0.02 | 2.24 | 76.06 |
| PO1_X5 | ML | XIXIMI-5 | LCs | 4.00 | 29.49 | 36.41 | 22.96 | 210.68 | 0.91 | 25.48523 | -88.00782 | 26.80 | 254.68 | Donwelling | 0.25 | 1.32 | 61.48 |
| TS1_X5 | ML | XIXIMI-5 | North | 9.00 | 28.09 | 34.44 | 21.95 | 218.03 | 1.19 | 20.52488 | -92.51078 | 18.90 | 65.56 | Upwelling | 0.03 | 1.38 | 81.11 |
| A10_X6 | ML | XIXIMI-6 | LCs | 3.00 | 30.26 | 36.30 | 22.61 | 188.98 | 0.12 | 24.94227 | -87.05583 | 41.50 | 216.53 | Donwelling | 0.00 | 1.50 | 100.55 |
| B17_X6 | ML | XIXIMI-6 | North | 3.00 | 29.91 | 36.21 | 22.67 | 189.99 | 0.12 | 24.00070 | -89.00077 | 27.70 | 102.87 | Neutral | 0.08 | 2.13 | 96.96 |
| C21_X6 | ML | XIXIMI-6 | Center | 3.00 | 30.53 | 36.55 | 22.71 | 189.77 | 0.11 | 22.99962 | -95.50135 | 46.50 | 112.76 | Neutral | 0.98 | 2.47 | 126.45 |
| C25_X6 | ML | XIXIMI-6 | Center | 3.00 | 29.76 | 35.95 | 22.52 | 186.07 | 0.16 | 22.99983 | -91.00292 | 27.70 | 116.20 | Neutral | 0.19 | 1.66 | 92.67 |
| D27_X6 | ML | XIXIMI-6 | Center | 3.00 | 30.18 | 36.52 | 22.80 | 190.62 | 0.11 | 22.00027 | -96.00145 | 27.70 | 103.38 | Neutral | 0.00 | 1.67 | 95.01 |
| F37_X6 | ML | XIXIMI-6 | Center | 3.00 | 29.83 | 36.56 | 22.96 | 191.68 | 0.15 | 21.00096 | -94.99710 | 28.70 | 103.38 | Neutral | 0.34 | 2.74 | 234.76 |
| G40_X6 | ML | XIXIMI-6 | South | 3.00 | 29.41 | 36.60 | 23.13 | 189.46 | 0.15 | 20.50140 | -96.00000 | 20.80 | 85.10 | Upwelling | 0.00 | 2.42 | 116.49 |
| G44_X6 | ML | XIXIMI-6 | South | 3.00 | 29.82 | 36.25 | 22.73 | 183.81 | 0.14 | 20.52263 | -93.50008 | 22.80 | 64.30 | Upwelling | 0.54 | 2.40 | 93.53 |
| H45_X6 | ML | XIXIMI-6 | South | 3.00 | 29.68 | 36.33 | 22.84 | 187.28 | 0.16 | 20.33333 | -95.60497 | 18.80 | 82.60 | Upwelling | 0.04 | 4.21 | 114.62 |
| Y2_X6 | ML | XIXIMI-6 | LCs | 5.00 | 29.53 | 35.87 | 22.60 | 187.00 | 0.10 | 21.62273 | -86.33987 | 29.90 | 116.50 | Neutral | 0.02 | 2.70 | 87.47 |
| Y3_X6 | ML | XIXIMI-6 | LCs | 3.00 | 29.41 | 36.16 | 22.80 | 192.39 | 0.15 | 21.64193 | -86.23207 | 18.80 | 120.17 | Neutral | 0.25 | 1.95 | 458.62 |
| Y7_X6 | ML | XIXIMI-6 | LCs | 3.00 | 30.28 | 36.19 | 22.53 | 190.53 | 0.13 | 21.65975 | -85.94695 | 33.60 | 154.28 | Donwelling | 0.90 | 2.85 | 85.98 |
| A10_X7 | ML | XIXIMI-7 | LCs | 5.00 | 28.33 | 36.24 | 23.22 | 201.34 | 0.00 | 24.99650 | -86.99704 | 52.65 | 238.32 | Donwelling | 0.04 | 1.57 | 96.87 |
| A2_X7 | ML | XIXIMI-7 | North | 9.00 | 27.53 | 36.43 | 23.63 | 203.79 | 0.10 | 24.87515 | -94.97753 | 25.83 | 151.47 | Neutral | 0.04 | 2.01 | 64.42 |
| A6_X7 | ML | XIXIMI-7 | North | 5.00 | 27.90 | 36.37 | 23.46 | 204.97 | 0.04 | 25.00122 | -91.00125 | 30.80 | 84.93 | Upwelling | 0.03 | 1.72 | 76.97 |
| A7_X7 | ML | XIXIMI-7 | North | 5.00 | 27.91 | 36.40 | 23.48 | 202.70 | 0.02 | 24.99887 | -89.99920 | 20.86 | 93.87 | Upwelling | 0.02 | 1.92 | 75.05 |
| B12_X7 | ML | XIXIMI-7 | North | 9.00 | 27.25 | 36.40 | 23.69 | 203.24 | 0.15 | 23.99867 | -95.06906 | 26.83 | 155.00 | Neutral | 0.04 | 2.10 | 74.24 |
| C23_X7 | ML | XIXIMI-7 | Center | 7.00 | 28.15 | 36.68 | 23.61 | 200.02 | 0.07 | 22.99850 | -93.00275 | 37.76 | 132.62 | Neutral | 0.03 | 1.14 | 100.12 |
| C24_X7 | ML | XIXIMI-7 | Center | 8.00 | 28.33 | 36.65 | 23.53 | 200.86 | 0.04 | 23.00305 | -91.99999 | 28.82 | 130.63 | Neutral | 0.02 | 1.26 | 87.20 |
| Y6_X7 | ML | XIXIMI-7 | LCs | 9.80 | 28.68 | 36.18 | 23.06 | 196.89 | 0.02 | 21.66085 | -86.06042 | 93.40 | 142.56 | Neutral | 0.08 | 1.53 | 71.49 |
| Y7_X7 | ML | XIXIMI-7 | LCs | 5.00 | 28.60 | 36.16 | 23.07 | 201.02 | 0.04 | 21.68740 | -85.93642 | 36.70 | 156.96 | Neutral | 0.04 | 1.51 | 76.16 |
| Y9_X7 | ML | XIXIMI-7 | LCs | 6.00 | 29.06 | 36.22 | 22.96 | 199.60 | 0.04 | 20.75658 | -85.61938 | 45.71 | 243.85 | Donwelling | 0.00 | 1.64 | 83.65 |

Temp=Temperature (°C), Abs. Salinity= Absolute salinity $S_A$(g. $kg^{-1}$), Fluor= fluorescence (relative fluorescence units (RFU)), MLD= Mixed layer depth(m), Nutri= Nutricline depth (m), Iso. 25.5= 25.5 kg $m^{-3}$ Isopycnal state, Nitrito= Nitrate+nitrite , Si= silicic acid, DOC= Disolved organic carbon
