## Supplementary material for "Response of microbial eukaryote community to the oligotrophic waters of the Gulf of Mexico: a plausible scenario for warm and stratified oceans": Table S5. ASVs co-occurrence associations from the ML

| Var1 | Var2 | Weight |
|---|---|---|
| ASV_00077:Chrysochromulina | ASV_00128:o__Prymnesiophyceae | 0.33330846 |
| ASV_00039:g__Gymnodiniaceae | ASV_00043:Scrippsiella | 0.31858961 |
| ASV_00131:MAST-3E_XX | ASV_00460:Haptophyta_Clade_HAP3_X | 0.28795113 |
| ASV_00170:c__Haptophyta | ASV_00239:f__Dictyochophyceae_X | 0.2168771 |
| ASV_00005:Gymnodinium | ASV_00006:o__Dinophyceae | 0.19341287 |
| ASV_00047:Karlodinium | ASV_00131:MAST-3E_XX | 0.16677231 |
| ASV_00308:Pedinellales_X | ASV_00403:MAST-7D_XX | 0.16379486 |
| ASV_00032:g__Prymnesiaceae | ASV_00128:o__Prymnesiophyceae | 0.16361541 |
| ASV_00058:g__Telonemia-Group-2 | ASV_00077:Chrysochromulina | 0.15916473 |
| ASV_00244:Pterocystida_XX | ASV_00298:Dictyochales_X | 0.15889172 |
| ASV_00104:Braarudosphaera | ASV_00244:Pterocystida_XX | 0.13131476 |
| ASV_00244:Pterocystida_XX | ASV_00407:Haptophyta_XXXX | 0.1283786 |
| ASV_00163:g__Raphid-pennate | ASV_00276:Torodinium | 0.11836368 |
| ASV_00006:o__Dinophyceae | ASV_00039:g__Gymnodiniaceae | 0.10976037 |
| ASV_00214:MAST-4C_XX | ASV_00421:MAST-11_XXX | 0.09877664 |
| ASV_00215:f__Diplonemea | ASV_00298:Dictyochales_X | 0.0960598 |
| ASV_00122:RAD-B-Group-IV_X | ASV_00212:RAD-A_XXX | 0.09052391 |
| ASV_00039:g__Gymnodiniaceae | ASV_00350:c__Ciliophora | 0.08949886 |
| ASV_00371:Dino-Group-I-Clade-4_X | ASV_00422:MAST-9A_XX | 0.08611008 |
| ASV_00128:o__Prymnesiophyceae | ASV_00357:Pterosperma | 0.07927464 |
| ASV_00244:Pterocystida_XX | ASV_00275:g__Arthracanthida-Symphya | 0.07634237 |
| ASV_00104:Braarudosphaera | ASV_00214:MAST-4C_XX | 0.07290951 |
| ASV_00298:Dictyochales_X | ASV_00319:Chrysophyceae_Clade-G_X | 0.06533212 |
| ASV_00039:g__Gymnodiniaceae | ASV_00625:f__Strombidiida | 0.06147143 |
| ASV_00246:MOCH-2_XXX | ASV_00403:MAST-7D_XX | 0.05883015 |
| ASV_00131:MAST-3E_XX | ASV_00163:g__Raphid-pennate | 0.05616769 |
| ASV_00296:Pentapharsodinium | ASV_00421:MAST-11_XXX | 0.0555021 |
| ASV_00296:Pentapharsodinium | ASV_00330:Goniomonas | 0.05397555 |
| ASV_00298:Dictyochales_X | ASV_00308:Pedinellales_X | 0.05332729 |
| ASV_00043:Scrippsiella | ASV_00169:Levanderina | 0.04338858 |
| ASV_00246:MOCH-2_XXX | ASV_00407:Haptophyta_XXXX | 0.03986402 |
| ASV_00244:Pterocystida_XX | ASV_00308:Pedinellales_X | 0.03929563 |
| ASV_00047:Karlodinium | ASV_00078:Picomonas | 0.03893099 |
| ASV_00189:Chaetoceros | ASV_00566:Partenskyella | 0.03430265 |
| ASV_00047:Karlodinium | ASV_00215:f__Diplonemea | 0.03213165 |
| ASV_00021:Gyrodinium | ASV_00047:Karlodinium | 0.03204904 |
| ASV_00112:g__Dino-Group-I-Clade-4 | ASV_00172:Hemiselmis | 0.03195652 |
| ASV_00272:MAST-7B_XX | ASV_00330:Goniomonas | 0.02788347 |
| ASV_00244:Pterocystida_XX | ASV_00451:MAST-3C_XX | 0.02782724 |
| ASV_00135:Azadinium | ASV_00481:f__Peridiniales | 0.02767655 |
| ASV_00103:RAD-B-Group-II_X | ASV_00122:RAD-B-Group-IV_X | 0.02624465 |
| ASV_00403:MAST-7D_XX | ASV_00407:Haptophyta_XXXX | 0.02475796 |
| ASV_00298:Dictyochales_X | ASV_00439:Chrysophyceae_Clade-H_X | 0.02447956 |
| ASV_00104:Braarudosphaera | ASV_00308:Pedinellales_X | 0.02191667 |
| ASV_00239:f__Dictyochophyceae_X | ASV_00439:Chrysophyceae_Clade-H_X | 0.02097039 |
| ASV_00319:Chrysophyceae_Clade-G_X | ASV_00403:MAST-7D_XX | 0.02051475 |

**Table S5. ASVs co-occurrence associations from the ML (continued)**

| Var1 | Var2 | Weight |
|---|---|---|
| ASV_00032:g__Prymnesiaceae | ASV_00050:MAST-1D_XX | 0.01826685 |
| ASV_00350:c__Ciliophora | ASV_00552:Strombidiida_G_XX | 0.00853743 |
| ASV_00039:g__Gymnodiniaceae | ASV_00203:Heterocapsa | 0.00834099 |
| ASV_00214:MAST-4C_XX | ASV_00298:Dictyochales_X | 0.00629588 |
| ASV_00005:Gymnodinium | ASV_00273:Alexandrium | 1.68E-04 |
| ASV_00112:g__Dino-Group-I-Clade-4 | ASV_00439:Chrysophyceae_Clade-H_X | -0.00575385 |
| ASV_00425:MAST-3I_XX | ASV_00686:Protoperidinium | -0.00791723 |
| ASV_00233:MAST-3A_XX | ASV_00625:f__Strombidiida | -0.00988428 |
| ASV_00099:Paragymnodinium | ASV_00215:f__Diplonemea | -0.01396766 |
| ASV_00112:g__Dino-Group-I-Clade-4 | ASV_00298:Dictyochales_X | -0.02359777 |
| ASV_00119:Blastodinium | ASV_00215:f__Diplonemea | -0.0313276 |
| ASV_00308:Pedinellales_X | ASV_00624:Pelagodinium | -0.03132958 |
| ASV_00308:Pedinellales_X | ASV_00416:f__Bacillariophyta_X | -0.03586561 |
| ASV_00425:MAST-3I_XX | ASV_00625:f__Strombidiida | -0.03912588 |
| ASV_00319:Chrysophyceae_Clade-G_X | ASV_00772:Chloropicon | -0.04550578 |
| ASV_00126:MAST-4B_XX | ASV_00772:Chloropicon | -0.0525384 |
| ASV_00298:Dictyochales_X | ASV_00339:Aggregata | -0.05367498 |
| ASV_00112:g__Dino-Group-I-Clade-4 | ASV_00403:MAST-7D_XX | -0.05907484 |
| ASV_00039:g__Gymnodiniaceae | ASV_00386:Margalefidinium | -0.07007834 |
| ASV_00119:Blastodinium | ASV_00298:Dictyochales_X | -0.07332477 |
| ASV_00122:RAD-B-Group-IV_X | ASV_00128:o__Prymnesiophyceae | -0.08291011 |
| ASV_00172:Hemiselmis | ASV_01091:Eutintinnus | -0.0866162 |
| ASV_00172:Hemiselmis | ASV_00439:Chrysophyceae_Clade-H_X | -0.10924289 |
| ASV_00006:o__Dinophyceae | ASV_00233:MAST-3A_XX | -0.1226748 |
| ASV_00403:MAST-7D_XX | ASV_00624:Pelagodinium | -0.1656126 |
| ASV_00039:g__Gymnodiniaceae | ASV_00425:MAST-3I_XX | -0.19302998 |
