## Supplementary material for "Response of microbial eukaryote community to the oligotrophic waters of the Gulf of Mexico: a plausible scenario for warm and stratified oceans": Table S6. ASVs co-occurrence associations from the DCM

| Var1 | Var2 | Weight |
|---|---|---|
| ASV_00058:g__Telonemia-Group-2 | ASV_00128:o__Prymnesiophyc | 0.3020943 |
| ASV_00045:Karlodinium | ASV_00685:g__Arthracanthida- | 0.27367751 |
| ASV_00058:g__Telonemia-Group-2 | ASV_00077:Chrysochromulina | 0.24673429 |
| ASV_00006:o__Dinophyceae | ASV_00021:Gyrodinium | 0.24616445 |
| ASV_00005:Gymnodinium | ASV_00006:o__Dinophyceae | 0.24543662 |
| ASV_00008:Pelagomonas | ASV_00107:Phaeocystis | 0.23244292 |
| ASV_00006:o__Dinophyceae | ASV_00786:f__Bacillariophyta_ | 0.2242356 |
| ASV_00232:f__Diplonemea | ASV_00641:DSPD-1_X | 0.22155554 |
| ASV_00101:Bathycoccus | ASV_00491:Dino-Group-II-Clade | 0.21194572 |
| ASV_00210:c__Ciliophora | ASV_00279:g__Dino-Group-I-Cl | 0.20855168 |
| ASV_00006:o__Dinophyceae | ASV_00804:Pseudocubus | 0.20458735 |
| ASV_00005:Gymnodinium | ASV_00054:f__Gymnodiniales | 0.1931759 |
| ASV_00050:MAST-1D_XX | ASV_00078:Picomonas | 0.16908785 |
| ASV_00005:Gymnodinium | ASV_00161:Torodinium | 0.16266696 |
| ASV_00045:Karlodinium | ASV_01121:MAST-3E_XX | 0.16124081 |
| ASV_00242:g__Chaunacanthida_X | ASV_00434:RAD-C_XXX | 0.15506745 |
| ASV_00242:g__Chaunacanthida_X | ASV_00418:g__Picozoa_XXX | 0.15113927 |
| ASV_00092:Dino-Group-II-Clade-22_X | ASV_00123:Ostreococcus | 0.14373035 |
| ASV_00021:Gyrodinium | ASV_00086:Lepidodinium | 0.14315647 |
| ASV_00154:Pseudo-nitzschia | ASV_00377:Thalassiosira | 0.14250037 |
| ASV_00119:Blastodinium | ASV_00539:o__Spirotrichea | 0.13242116 |
| ASV_00058:g__Telonemia-Group-2 | ASV_00349:MOCH-4_XXX | 0.1245941 |
| ASV_00065:Dino-Group-I-Clade-1_X | ASV_00184:c__Dinoflagellata | 0.11271884 |
| ASV_00120:g__Spongodiscidae-Coccodiscida | ASV_00635:f__Spumellarida | 0.11128234 |
| ASV_00336:f__Dino-Group-II | ASV_00358:Dino-Group-II-Clade | 0.10904139 |
| ASV_00170:c__Haptophyta | ASV_00756:o__Acantharea | 0.10613278 |
| ASV_00210:c__Ciliophora | ASV_00337:MAST-3I_XX | 0.09880863 |
| ASV_00107:Phaeocystis | ASV_00132:Prymnesiophyceae_ | 0.0936615 |
| ASV_00054:f__Gymnodiniales | ASV_00786:f__Bacillariophyta_ | 0.09171238 |
| ASV_00158:g__RAD-B-Group-IV | ASV_00491:Dino-Group-II-Clade | 0.08371408 |
| ASV_00110:MAST-1C_XX | ASV_00210:c__Ciliophora | 0.07927976 |
| ASV_00065:Dino-Group-I-Clade-1_X | ASV_00250:g__Sphaerozoidae | 0.07097083 |
| ASV_00283:g__Cryptomonadales_X | ASV_00810:MAST-3D_XX | 0.06588364 |
| ASV_00150:RAD-B-Group-I_X | ASV_00345:Dino-Group-II-Clade | 0.05782328 |
| ASV_00008:Pelagomonas | ASV_00132:Prymnesiophyceae_ | 0.05383178 |
| ASV_00409:Nassellaria_XX | ASV_00804:Pseudocubus | 0.0488083 |
| ASV_00336:f__Dino-Group-II | ASV_00448:Dino-Group-V_XX | 0.04578834 |
| ASV_00135:Azadinium | ASV_00483:f__Peridiniales | 0.04356735 |
| ASV_00073:RAD-A_XXX | ASV_00128:o__Prymnesiophyc | 0.04100899 |
| ASV_00218:c__Chlorophyta | ASV_00395:Chloropicon | 0.03753021 |
| ASV_00092:Dino-Group-II-Clade-22_X | ASV_00232:f__Diplonemea | 0.03662454 |
| ASV_00150:RAD-B-Group-I_X | ASV_00333:RAD-B_XXX | 0.03618106 |
| ASV_00150:RAD-B-Group-I_X | ASV_00474:Dino-Group-II-Clade | 0.03454136 |
| ASV_00127:f__Dinophysiales | ASV_00135:Azadinium | 0.03241695 |
| ASV_00045:Karlodinium | ASV_00078:Picomonas | 0.02912167 |
| ASV_00154:Pseudo-nitzschia | ASV_00777:Chaetoceros | 0.02349352 |
| ASV_00065:Dino-Group-I-Clade-1_X | ASV_00924:Dino-Group-I-Clade | 0.02345666 |

**Table S6. ASVs co-occurrence associations from the DCM (continued)**

| Var1 | Var2 | Weight |
|---|---|---|
| ASV_00161:Torodinium | ASV_00184:c__Dinoflagellata | 0.02321107 |
| ASV_00006:o__Dinophyceae | ASV_00161:Torodinium | 0.02116481 |
| ASV_00248:MOCH-2_XXX | ASV_00337:MAST-3I_XX | 0.01811176 |
| ASV_00377:Thalassiosira | ASV_00590:g__Polar-centric-M | 0.01750074 |
| ASV_00123:Ostreococcus | ASV_00590:g__Polar-centric-M | 0.01642778 |
| ASV_00032:g__Prymnesiaceae | ASV_00170:c__Haptophyta | 0.0082379 |
| ASV_00248:MOCH-2_XXX | ASV_00530:Syndiniales_XXX | 0.00664508 |
| ASV_00382:Prymnesiophyceae_Clade_B3_X | ASV_01024:Protoperidinium | 0.00367547 |
| ASV_00119:Blastodinium | ASV_00279:g__Dino-Group-I-Cl | 0.00149982 |
| ASV_00138:Gephyrocapsa | ASV_00434:RAD-C_XXX | -2.45E-04 |
| ASV_00279:g__Dino-Group-I-Clade-4 | ASV_00491:Dino-Group-II-Clade | -0.00277365 |
| ASV_00050:MAST-1D_XX | ASV_00107:Phaeocystis | -0.00819016 |
| ASV_00073:RAD-A_XXX | ASV_00198:g__Raphid-pennate | -0.01324726 |
| ASV_00073:RAD-A_XXX | ASV_00786:f__Bacillariophyta_ | -0.01363597 |
| ASV_00135:Azadinium | ASV_00234:RAD-B-Group-IV_X | -0.01499926 |
| ASV_00239:f__Dictyochophyceae_X | ASV_00539:o__Spirotrichea | -0.01503298 |
| ASV_00092:Dino-Group-II-Clade-22_X | ASV_00138:Gephyrocapsa | -0.01550397 |
| ASV_00234:RAD-B-Group-IV_X | ASV_00283:g__Cryptomonadal | -0.01599433 |
| ASV_00170:c__Haptophyta | ASV_00924:Dino-Group-I-Clade | -0.01826554 |
| ASV_00008:Pelagomonas | ASV_00685:g__Arthracanthida- | -0.01952432 |
| ASV_00086:Lepidodinium | ASV_00240:f__RAD-B_X | -0.02002253 |
| ASV_00337:MAST-3I_XX | ASV_00480:MAST-4C_XX | -0.02357852 |
| ASV_00078:Picomonas | ASV_00107:Phaeocystis | -0.02744866 |
| ASV_00033:g__Spumellarida-Group-I | ASV_00786:f__Bacillariophyta_ | -0.04670079 |
| ASV_00008:Pelagomonas | ASV_00078:Picomonas | -0.05317222 |
| ASV_00101:Bathycoccus | ASV_00279:g__Dino-Group-I-Cl | -0.05692678 |
| ASV_00198:g__Raphid-pennate | ASV_00337:MAST-3I_XX | -0.0593364 |
| ASV_00158:g__RAD-B-Group-IV | ASV_00530:Syndiniales_XXX | -0.08799052 |
| ASV_00021:Gyrodinium | ASV_00756:o__Acantharea | -0.09055299 |
| ASV_00418:g__Picozoa_XXX | ASV_00985:Alexandrium | -0.09246324 |
| ASV_00345:Dino-Group-II-Clade-7_X | ASV_00539:o__Spirotrichea | -0.09399173 |
| ASV_00039:g__Gymnodiniaceae | ASV_00242:g__Chaunacanthida | -0.11720474 |
| ASV_00050:MAST-1D_XX | ASV_00161:Torodinium | -0.14130591 |
| ASV_00099:Paragymnodinium | ASV_00924:Dino-Group-I-Clade | -0.16419936 |
| ASV_00006:o__Dinophyceae | ASV_00242:g__Chaunacanthida | -0.19705314 |
