## Supplementary figures and statistic tabless. for "Response of microbial eukaryote community to the oligotrophic waters of the Gulf of Mexico: a plausible scenario for warm and stratified oceans"

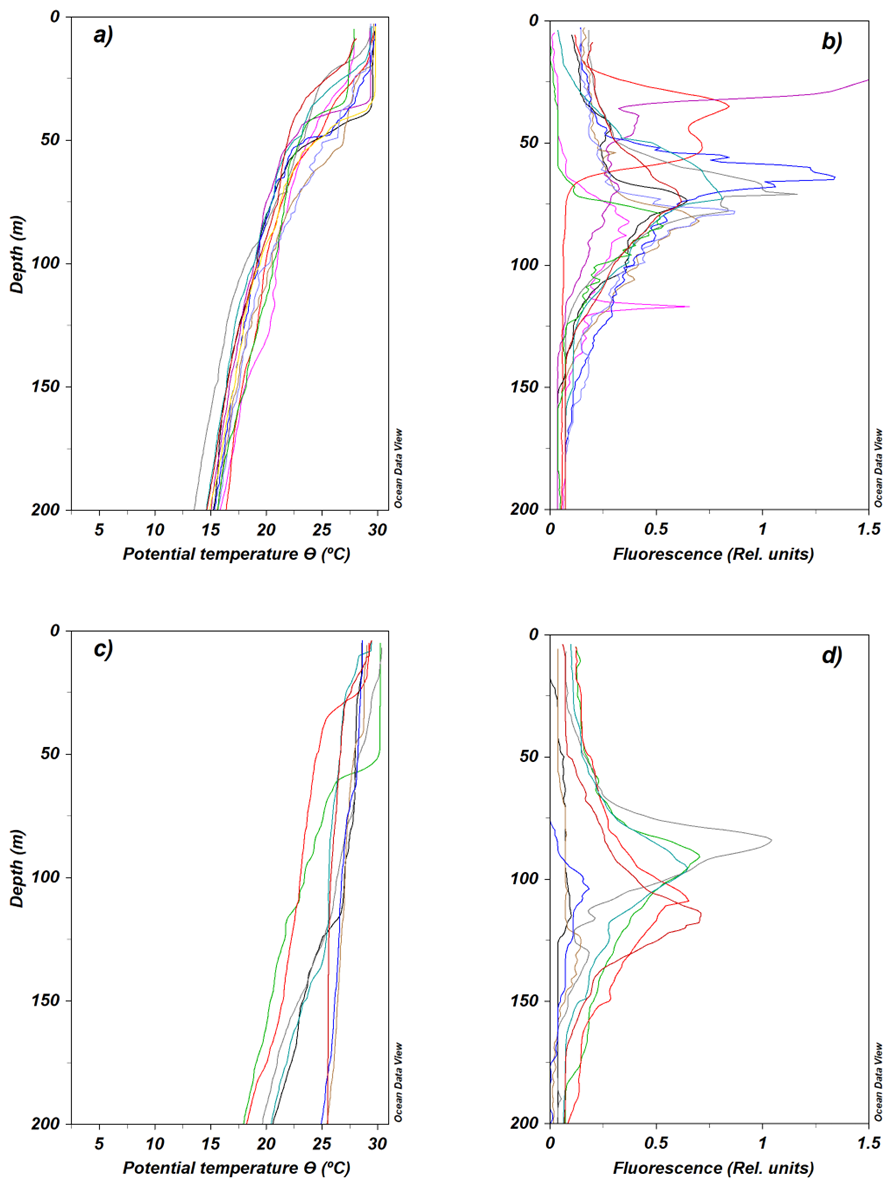

**Figure S1**. CTD profiles of potential temperature (ϴ; °C; *left)* and fluorescence (RFU *right*) under cyclonic conditions (a and b) compared to anticyclonic conditions (c and d).

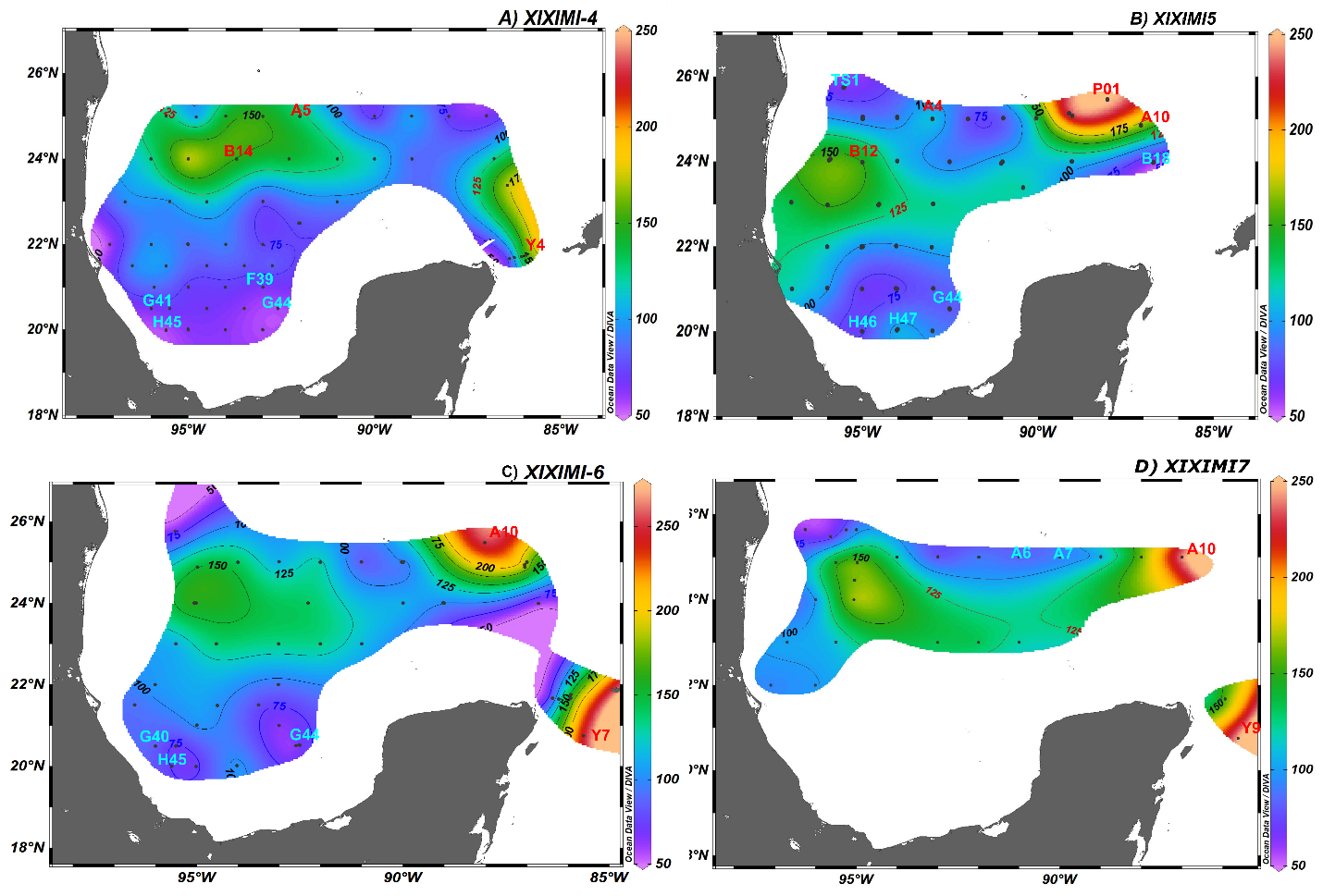

**Figure S2**. Vertical distribution of the subsurface 25.5 kg m^-3^ isopycnal. The isoline represents depth (m). The stations in blue are under cyclonic influence, and the stations in red are under anticyclonic influence.

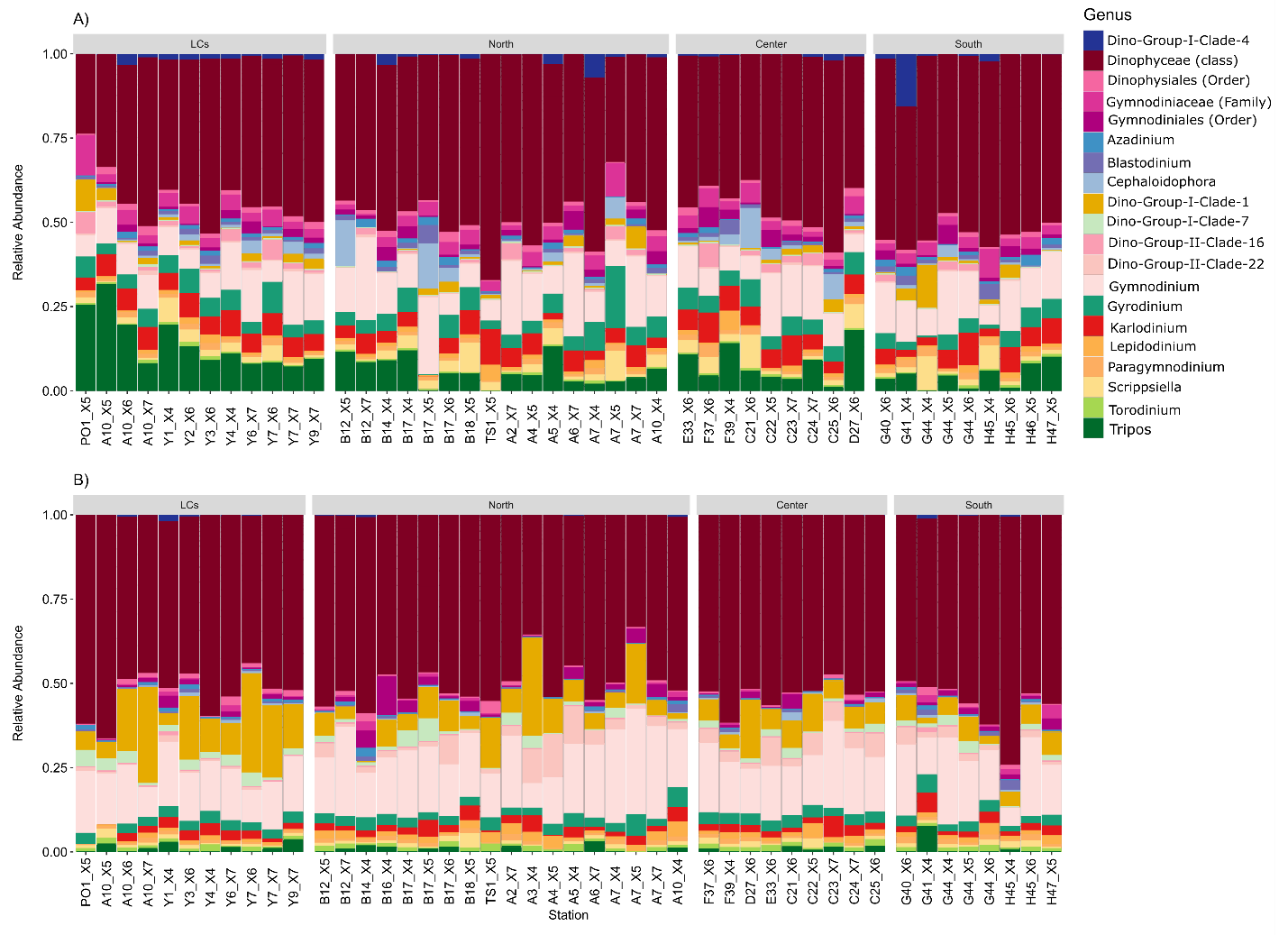
**Figure S3.** Relative read abundance of principal ASVs of Alveolata genus from A) mixed layer (ML) and B) depth chlorophyll maximum (DCM) from the regions of Loop Current (LCs, stations within the LC and Yucatan Channel), northern (region to the north of the Exclusive Economic Zone of Mexico (EEZ)), central (center of the Gulf of Mexico (GoM)), and southern (Bay of Campeche).

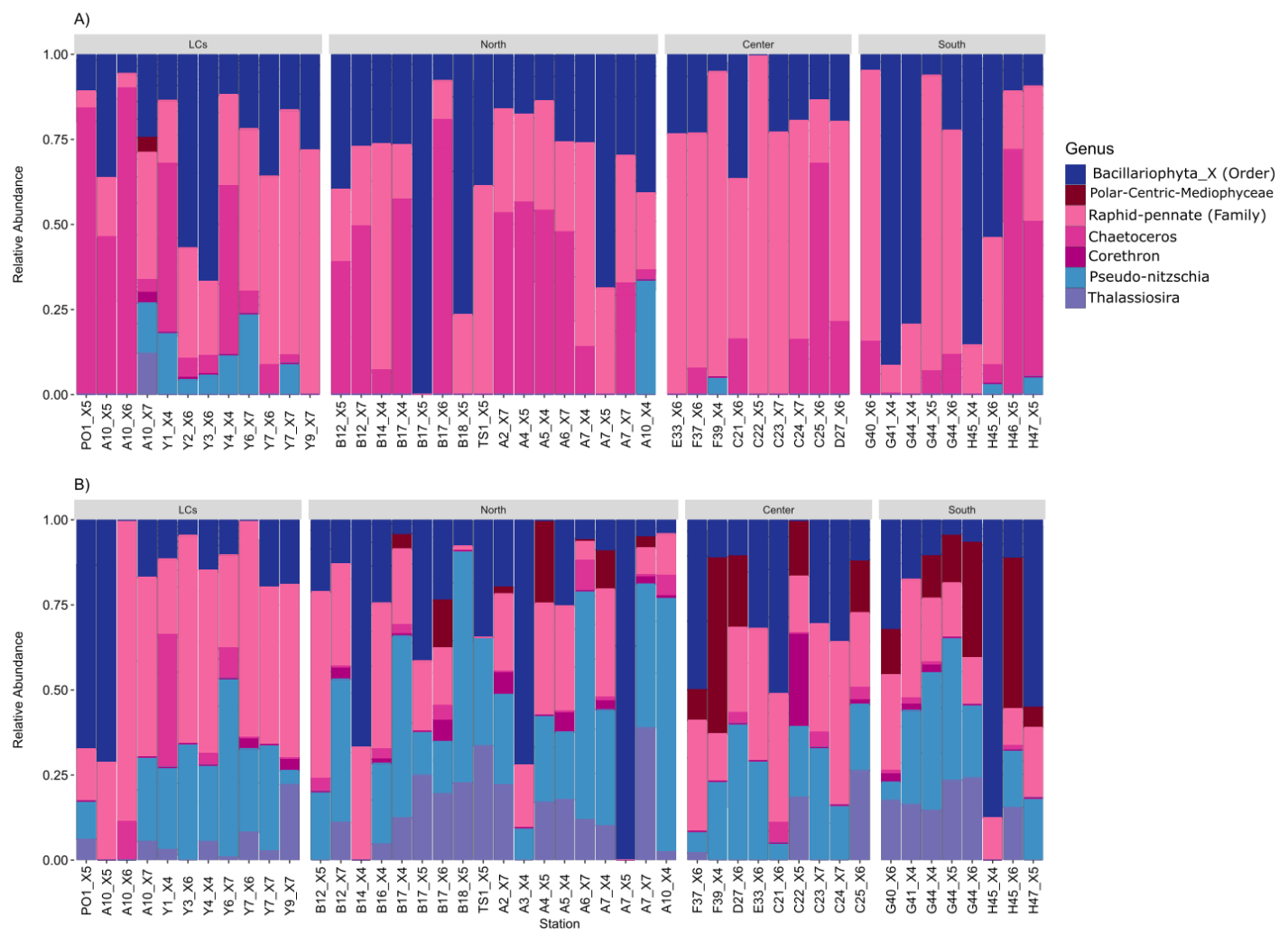
**Figure S4.** Relative read abundance of principal ASVs of Diatoms genus from A) mixed layer (ML) and B) depth chlorophyll maximum (DCM) from the regions of Loop Current (LCs, stations within the LC and Yucatan Channel), northern (region to the north of the Exclusive Economic Zone of Mexico (EEZ)), central (center of the Gulf of Mexico (GoM)), and southern (Bay of Campeche).

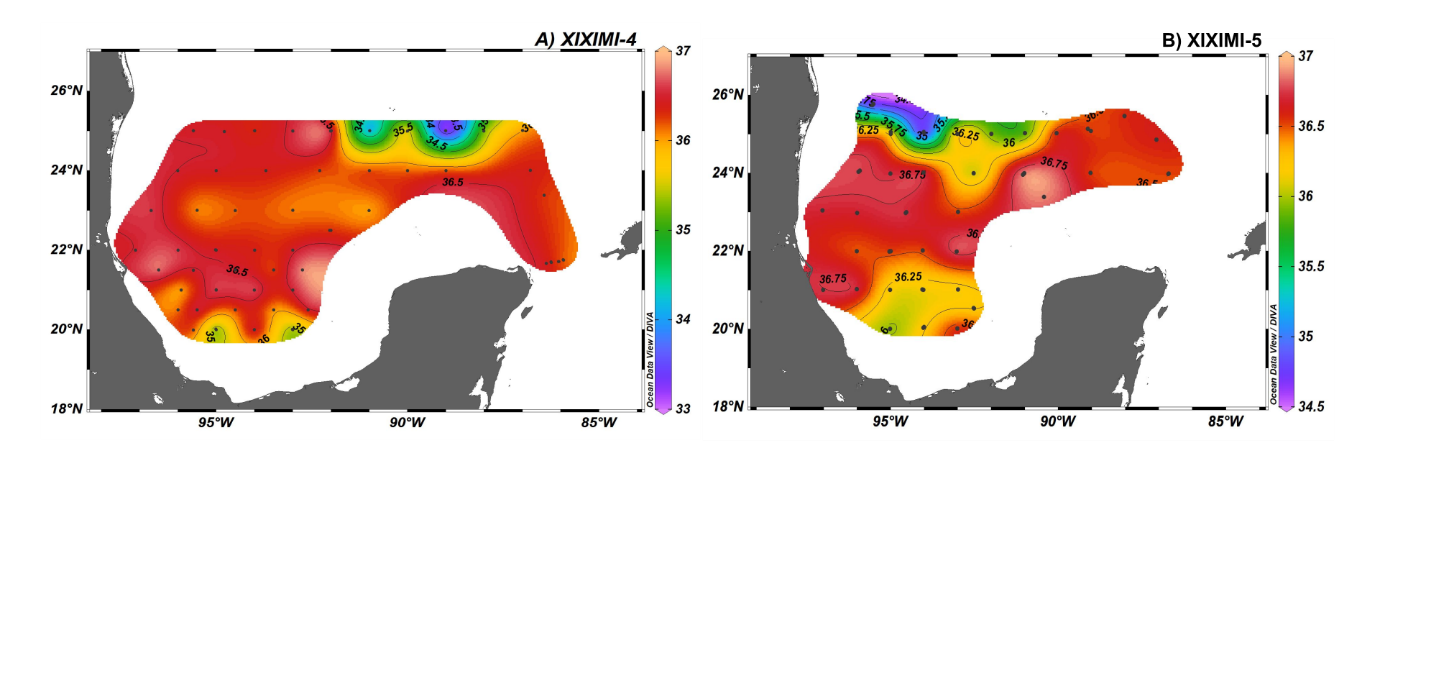

**Figure S5.** Spatial distribution of absolute salinity (S_A_ g kg^-1^) from the cruises XIXIMI-4 (2015) and XIXIMI-5 (2016).

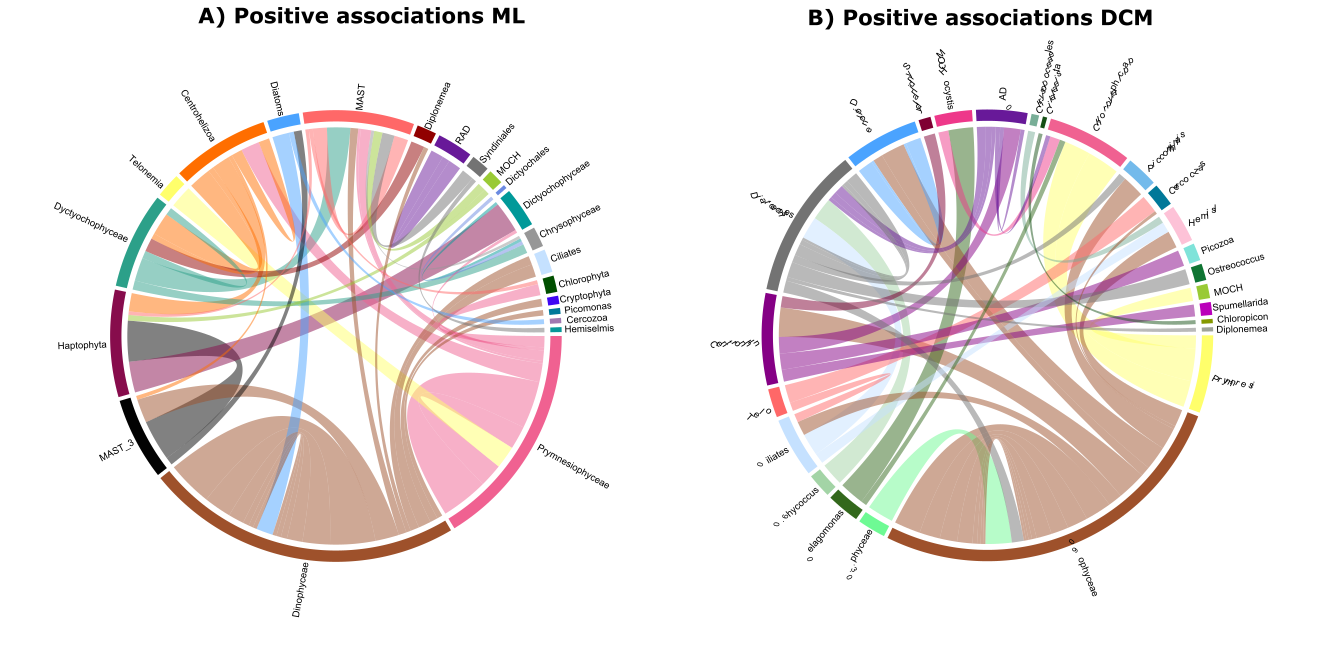

**Figure S6.** Protist ASVs Interaction network between protist-protist amplicon sequence variants (ASVs). Positive associations in the (A) mixed layer (ML) and (B) deep chlorophyll maximum (DCM). Ribbons represent the proportions of nodes (ASVs).

**Table S1.** Pairwise PERMANOVA comparison from the categorical variable A) cruise, B) region and C) nutricline depth from the ML (999 permutations).

| A) **Cruise comparison** | **R^2^** | **p.value** | **p.adjusted** |
| --- | --- | --- | --- |
| XIXIMI-5 vs XIXIMI-6 | 0.10 | 0.001 | **0.006*** |
| XIXIMI-5 vs XIXIMI-4 | 0.12 | 0.001 | **0.006*** |
| XIXIMI-5 vs XIXIMI-7 | 0.15 | 0.001 | **0.006*** |
| XIXIMI-6 vs XIXIMI-4 | 0.07 | 0.024 | 0.144 |
| XIXIMI-6 vs XIXIMI-7 | 0.16 | 0.001 | **0.006*** |
| XIXIMI-4 vs XIXIMI-7 | 0.21 | 0.001 | **0.006*** |
| B) **Region comparison** | **R^2^** | **p.value** | **p.adjusted** |
| LCs vs North | 0.08 | 0.001 | **0.006*** |
| LCs vs Center | 0.10 | 0.001 | **0.006*** |
| LCs vs South | 0.13 | 0.001 | **0.006*** |
| North vs Center | 0.05 | 0.298 | 1 |
| North vs South | 0.06 | 0.092 | 0.552 |
| Center vs South | 0.08 | 0.249 | 1 |
| C) **Nutricline depth comparison** | **R^2^** | **p.value** | **p.adjusted** |
| Downwelling vs Neutral | 0.04 | 0.234 | 1 |
| Downwelling vs Upwelling | 0.05 | 0.219 | 1 |
| Downwelling vs | 0.09 | 0.517 | 1 |
| Neutral vs Upwelling | 0.02 | 0.904 | 1 |
| Neutral vs | 0.05 | 0.547 | 1 |
| Upwelling vs | 0.07 | 0.575 | 1 |

**Table S2.** Pairwise PERMANOVA comparison from the categorical variable A) cruise, B) Region and C) Nutricline depth from the DCM (999 permutations).

| A) **Cruise comparison** | **R^2^** | **p.value** | **p.adjusted** |
| --- | --- | --- | --- |
| XIXIMI-5 vs XIXIMI-6 | 0.09 | 0.001 | **0.006*** |
| XIXIMI-5 vs XIXIMI-7 | 0.07 | 0.092 | 0.552 |
| XIXIMI-5 vs XIXIMI-4 | 0.08 | 0.006 | **0.036.** |
| XIXIMI-6 vs XIXIMI-7 | 0.08 | 0.025 | 0.15 |
| XIXIMI-6 vs XIXIMI-4 | 0.06 | 0.093 | 0.558 |
| XIXIMI-7 vs XIXIMI-4 | 0.10 | 0.014 | 0.084 |
| B) **Region comparison** | R^2^ | p.value | p.adjusted |
| Center vs LCs | 0.19 | 0.001 | **0.006*** |
| Center vs North | 0.05 | 0.045 | 0.27 |
| Center vs South | 0.10 | 0.034 | 0.204 |
| LCs vs North | 0.10 | 0.001 | **0.006*** |
| LCs vs South | 0.20 | 0.001 | **0.006*** |
| North vs South | 0.05 | 0.377 | 1 |
| C) **Nutricline depth comparison** | **R2** | **p.value** | **p.adjusted** |
| Neutral vs Downwelling | 0.07 | 0.001 | **0.003*** |
| Neutral vs Upwelling | 0.07 | 0.001 | **0.003*** |
| Downwelling vs Upwelling | 0.13 | 0.001 | **0.003*** |

**Table S3**. ANOVA analysis of betadisper by depth of the a) mixed layer (ML) and b) deep chlorphyll maximum (DCM) by categorical variables.

| a) **ANOVA BETADISPER MLD** | | | | |
| --- | --- | --- | --- | --- |
|  | **df** | **R^2^** | **F-value** | **p-value** |
| Cruise | 3 |  | 1.4388 | 0.23 |
| Region | 3 |  | 1.7604 | 0.173 |
| Nutricline depth | 2 |  | 1.6954 | 0.183 |
| b) **ANOVA BETADISPER DCM** | | | | |
|  | **df** | **R^2^** | **F-value** | **p-value** |
| Cruise | 3 |  | 3.0909 | **0.035** |
| Region | 3 |  | 3.9424 | **0.016** |
| Nutricline depth | 2 |  | 0.1568 | 0.85 |
